# Coregulation of extracellular vesicle production and fluconazole susceptibility in *Cryptococcus neoformans*

**DOI:** 10.1101/2023.01.30.526212

**Authors:** Juliana Rizzo, Adèle Trottier, Frédérique Moyrand, Jean-Yves Coppee, Corinne Maufrais, Ana Claudia G. Zimbres, Thi Tuong Vi Dang, Alexandre Alanio, Marie Desnos-Ollivier, Isabelle Mouyna, Gérard Péhau-Arnaudet, Pierre-Henri Commere, Sophie Novault, Iuliana V. Ene, Leonardo Nimrichter, Marcio L. Rodrigues, Guilhem Janbon

## Abstract

Resistance to fluconazole (FLC), the most widely used antifungal drug, is typically achieved by altering the azole drug target and/or drug efflux pumps. Recent reports have suggested a link between vesicular trafficking and antifungal resistance. Here, we identified novel *Cryptococcus neoformans* regulators of extracellular vesicle (EV) biogenesis that impact FLC resistance. In particular, the transcription factor Hap2 does not affect the expression of the drug target or efflux pumps, yet it impacts the cellular sterol profile. Subinhibitory FLC concentrations also downregulate EV production. Moreover, *in vitro* spontaneous FLC-resistant colonies showed altered EV production, and the acquisition of FLC resistance was associated with decreased EV production in clinical isolates. Finally, the reversion of FLC resistance was associated with increased EV production. These data suggest a model in which fungal cells can regulate EV production in place of regulating the drug target gene expression as a first line of defense against antifungal assault in this fungal pathogen.

**IMPORTANCE:** Extracellular vesicles (EVs) are membrane-enveloped particles that are released by cells into the extracellular space. Fungal EVs can mediate community interactions and biofilm formation but thier functions remain poorly understood. Here, we report the identification of the first regulators of EV production in the major fungal pathogen *Cryptococcus neoformans.* Surprisingly, we uncover a novel role of EVs in modulating antifungal drug resistance. Disruption of EV production was associated with altered lipid composition and changes in fluconazole susceptibility. Spontaneous azole-resistant mutants were deficient in EV production, while loss of resistance restored initial EV production levels. These findings were recapitulated in *C. neoformans* clinical isolates, indicating that azole resistance and EV production are coregulated in diverse strains. Our study reveals a new mechanism of drug resistance in which cells adapt to azole stress by modulating EV production.

## 1) INTRODUCTION

Fungal diseases have been consistently neglected despite their increasing impact on human health. These eukaryotic pathogens infect ∼1.5 billion people worldwide and lead to ∼1.5 million deaths annually^1, 2^. Over 90% of reported fungal-associated deaths are caused by species from four genera *Aspergillus*, *Cryptococcus*, *Candida*, or *Pneumocystis*^3^. The species *Cryptococcus neoformans*, *Candida albicans*, *Candida auris,* and *Aspergillus fumigatus* form the critical priority group within the fungal pathogens list published by the World Health Organization (WHO) in 2022^4, 5^. Fungal infections are generally difficult to treat, and mortality rates remain high despite available antifungal treatments^6^. The arsenal of antifungal molecules is based on four main classes: polyenes (e.g., amphotericin B), azoles (e.g., fluconazole (FLC)), echinocandins (e.g., caspofungin), and flucytosine (a pyrimidine analog)^7^. Due to their bioavailability, low toxicity, and wide spectrum of action, azoles are the most widely used antifungals. Azoles are fungistatic drugs that inhibit the cytochrome P450-dependent enzyme 14-α demethylase encoded by *ERG11* in fungi, thus interrupting the synthesis of ergosterol^6^. In azole-treated cells, an accumulation of toxic intermediate sterols is observed, increasing membrane permeability and inhibiting fungal growth^8^. Prolonged use of FLC has the potential to select FLC-resistant strains^8^. Azole resistance has been associated with mutations in the *ERG11* sequence, thus limiting azole binding^8^, or in transcription factors (TF) like *TAC1* or *UPC1* regulating the expression of *ERG11* and/or drug efflux pumps^9, 10^. Heteroresistance has also been described in *C. neoformans*, and was associated with the unstable duplication of chromosome 1, resulting in the potential overexpression of *ERG11* and the efflux pump encoding gene *AFR1*^11, 12^.

Extracellular vesicles (EVs) are cell-derived membrane particles released to the extracellular space known to be produced in all domains of life. In fungi, they have been isolated from at least fifteen different genera^13^. *Cryptococcus* EVs display extensive diversity in size, shape, and structure^14^ and most EVs appear decorated by fibrillary material composed of mannoproteins, and are covered by capsule polysaccharide-like material^14^. EVs contain lipids (including ergosterol), polysaccharides, small molecules, pigments, and RNAs, although our understanding of the exact composition and function of these cargo molecules remains limited^12^.

The involvement of vesicular trafficking in fungal FLC resistance has been previously suggested. For instance, turbinmicin, a molecule with a broad spectrum antifungal activity, targets Sec14p in the post-Golgi vesicular trafficking pathway and synergizes with FLC against *C. albicans* biofilm formation^15^. Sortin2 inhibition of vesicular transport potentiates azoles in *C. albicans* and *C. glabrata*^16^. Studies in *C. neoformans* and *S. cerevisiae* have implicated Golgi trafficking and vesicle formation as potential avenues of azole potentiation^17–19^. Recent studies also suggested that EVs released by fungi could play a role in antifungal resistance^20–23^. For instance, *C. albicans* EVs confer FLC resistance to biofilms and ESCRT (endosomal sorting complexes required for transport) mutants defective in EVs production show altered biofilm FLC resistance^20^. EVs from *C. auris* drug-resistant strain induced amphotericin-B resistance to a susceptible strain^21^, but the mechanisms underlying this process remain largely unknown. Other examples of EVs associated with drug resistance come from the study of *Leishmania* parasites^24^, where EVs mediate the delivery of drug-resistance genes, leading to the emergence of resistant subpopulations^24^.

Despite these intriguing connections, the study of fungal EV biosynthesis in the context of antifungal resistance has been constrained by our limited understanding of EV biosynthesis. Genetic analyses of EV production and regulation have been hampered by the long and cumbersome protocols necessary to isolate and study these particles. Only a handful of mutations associated with defects in EV production have been reported so far, as recently reviewed^13^.

To narrow this knowedge gap, we screened a *C. neoformans* TF mutant library^25^ and identified the first regulators of EV biogenesis in fungi. We present data revealing a phenotypic association between drug resistance and regulation of EV biogenesis in these mutant strains but also in spontaneous FLC-resistant mutants isolated *in vitro,* as well as in clinical isolates. Moreover, we demonstrate that EV-defective mutants show altered lipid content. Finally, we show that sub-inhibitory concentrations of FLC regulate EV production. Taken together, our study points to a new mechanism of drug resistance in which cells adapt to FLC stress by modulating EV production. Our results uncover key regulators of EV biogenesis in *C. neoformans*, as well as a novel link with antifungal drug resistance.

## 2) RESULTS

### a) EV production is regulated by growth conditions and growth phase in *C. neoformans*

To test whether EV production varies across different media, growth stages, and temperatures, we optimized a new EV isolation protocol, in which the ultracentrifugation step was bypassed (**Fig 1A**). We used this protocol to demonstrate that EV production, as measured by the amount of total sterol in cellular supernatants, was highest when cells were grown in synthetic dextrose (SD) medium at 30°C compared to the other conditions tested (**Fig 1B**). By exploring the dynamics of EV production during the growth on SD agar plates at 30°C, we noticed that cells release EVs during a limited time window (16 to 24 h) corresponding to the transition between the exponential to stationary growth phase (**Fig 1C**).

**Figure 1:**
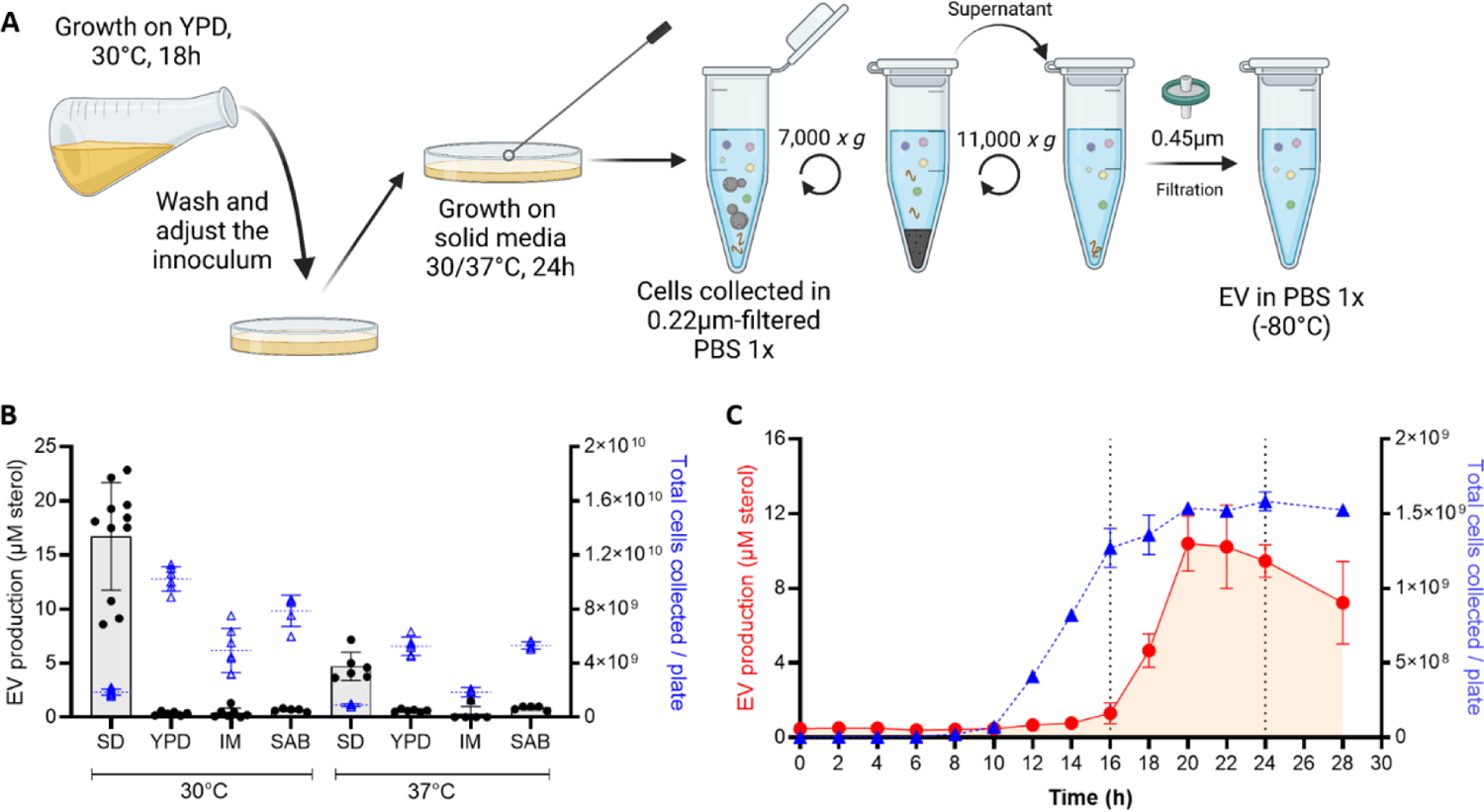
EV production is regulated by growth conditions and growth phase. Schematic representation of the optimized quick protocol used to obtain EVs from cells grown on solid medium (A). EV production from cells grown on different media: synthetic dextrose (SD), yeast extract-peptone-dextrose (YPD), capsule induction media (IM), and Sabouraud (SAB) at 30°C and 37°C, sterol concentration values are expressed per 10^9^ cells for each condition (B). EV production on SD medium at 30°C during the growth curve was analysed by the amount of total sterol in cellular supernatants when EVs are isolated using the quick protocol, values are expressed per plate of culture during each timepoint (C). Experiments were done with three or more biological replicates. Error bars show means ± SD. Schematic representation created in BioRender.

### b) Identification of regulators of EV production in *C. neoformans*

The tight regulation between EV production and growth suggested that specific TFs might regulate their biosynthesis. Thus, to identify potential regulators of EV biogenesis, we screened a collection of 155 TF mutants^25^ using the EV purification optimized protocol adjusted to 96-well plates (**Fig 2A**). We identified four mutants strongly altered in EV production. These strains produced <10% of the WT EV levels as evaluated by the total amount of sterols in EV samples. The EV-deficient strains lacked TFs *BZP2* (CNAG_04263), *GAT5* (CNAG_05153), *LIV4* (CNAG_06283) and *HAP2* (CNAG_07435). These results were confirmed when samples were prepared using the conventional untracentrifugation protocol^14^, measuring the total amount of sterols (**Fig 2B**), and evaluating the number of particles by nanoparticle flow cytometry (nanoFCM) analysis (**Fig 2C**). It is important to note that despite an impressive decrease in EV levels, these TF mutants were not completely impaired in EV production, as the high sensitivity of the nanoFCM analysis used detected EV-like particles in each of the EV samples obtained from the mutants. Moreover, using a large number of culture plates, we obtained enough *hap2Δ-*derived EVs to be observed by Cryo-EM. Image analysis revealed a similar structure of *hap2Δ* EVs as previously observed for WT EVs^14^ (**Fig 2D**).

**Figure 2:**
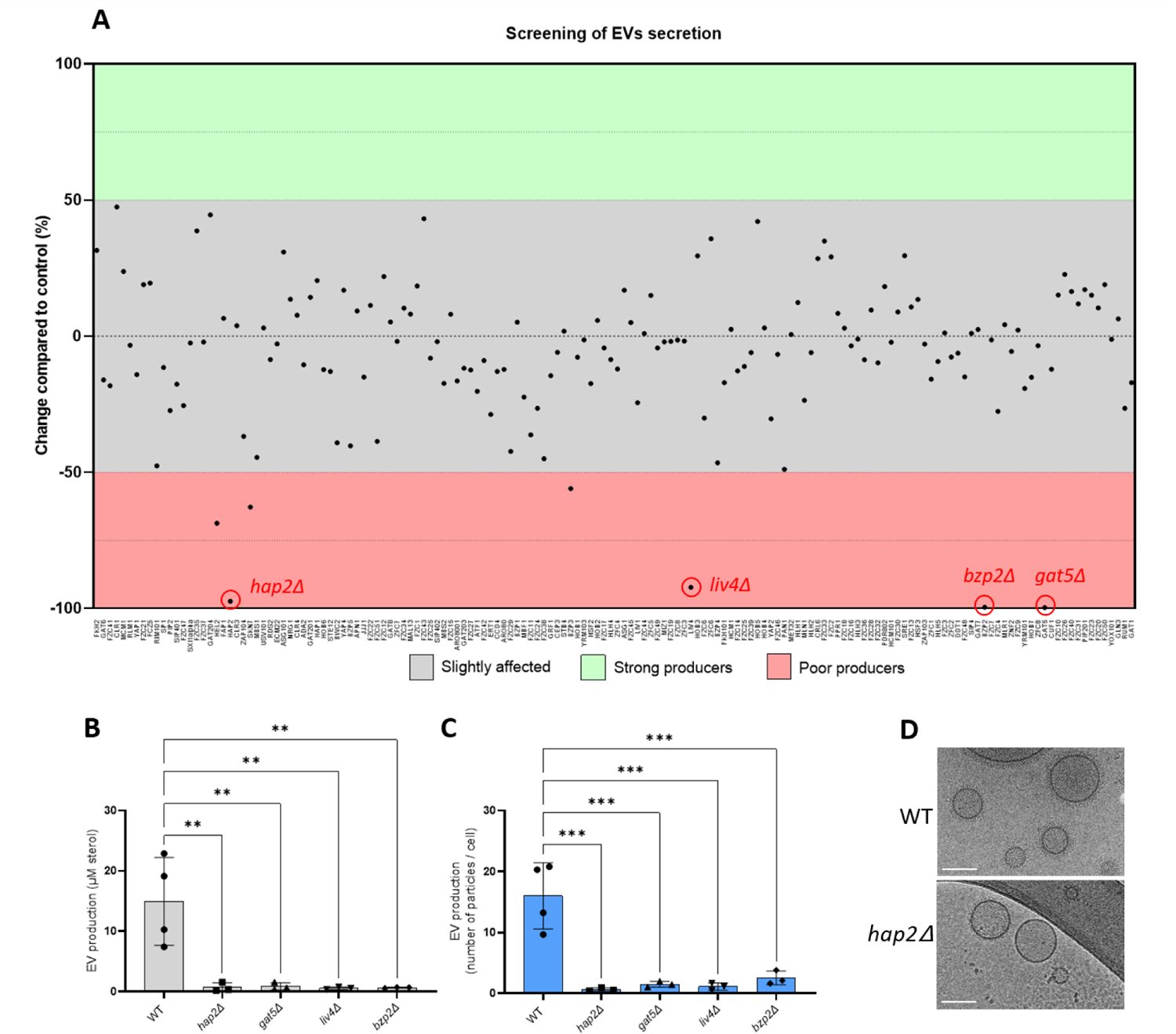
Identification of the transcription factors regulating EV production in *C. neoformans*. The screening of a 155 transcription factor (TF) mutant collection identified four major EV production regulators: *HAP2*, *LIV4*, *BZP2,* and *GAT5* (A). Confirmation of the EV phenotype using independent knockout strains generated in the KN99α background by measuring the amount of total sterol contained in EV samples. Sterol concentration values are expressed per 10^9^ cells for each condition(B) and single particle counting using nanoFCM (C). Cryo-EM analysis of EVs obtained from WT and *hap2Δ* strains (D), scale bar = 100 nm. Experiments were performed in at least three biological replicates. Error bars show means ± SD.

Bzp2p and Gat5p belong to a conserved zinc finger family containing TFs known as GATA-factors, regulating radiation sensitivity and nitrogen catabolite repression (NCR) when preferred nitrogen sources are absent or limiting^26^. Liv4p is a conserved MYB-like DNA-binding domain TF regulating growth and virulence in *C. neoformans*^27^. Hap2p is a subunit of the evolutionary conserved CCAAT-binding <u>h</u>eme <u>a</u>ctivator <u>p</u>rotein (HAP) complex, a heterotrimeric complex composed of Hap2/3/5 and the transcriptional activation subunit, HapX^28^. As expected, the *hap3Δ* and *hap5Δ* mutants were also drastically impaired for EV production. In contrast, the *hapXΔ* mutant strain produced the same quantity of EVs as the WT strain (**Fig 3A**), suggesting that the Hap2/3/5 complex controls EV production in an HapX-independent manner in *C. neoformans.* Notably, whereas *bzp2Δ*, *liv4Δ,* and *gat5Δ* mutant strains displayed reduced growth (**Fig S1**), none of the *hap* mutant strains showed growth deficiency under the conditions used to obtain EVs (**Fig 3B**).

**Figure 3:**
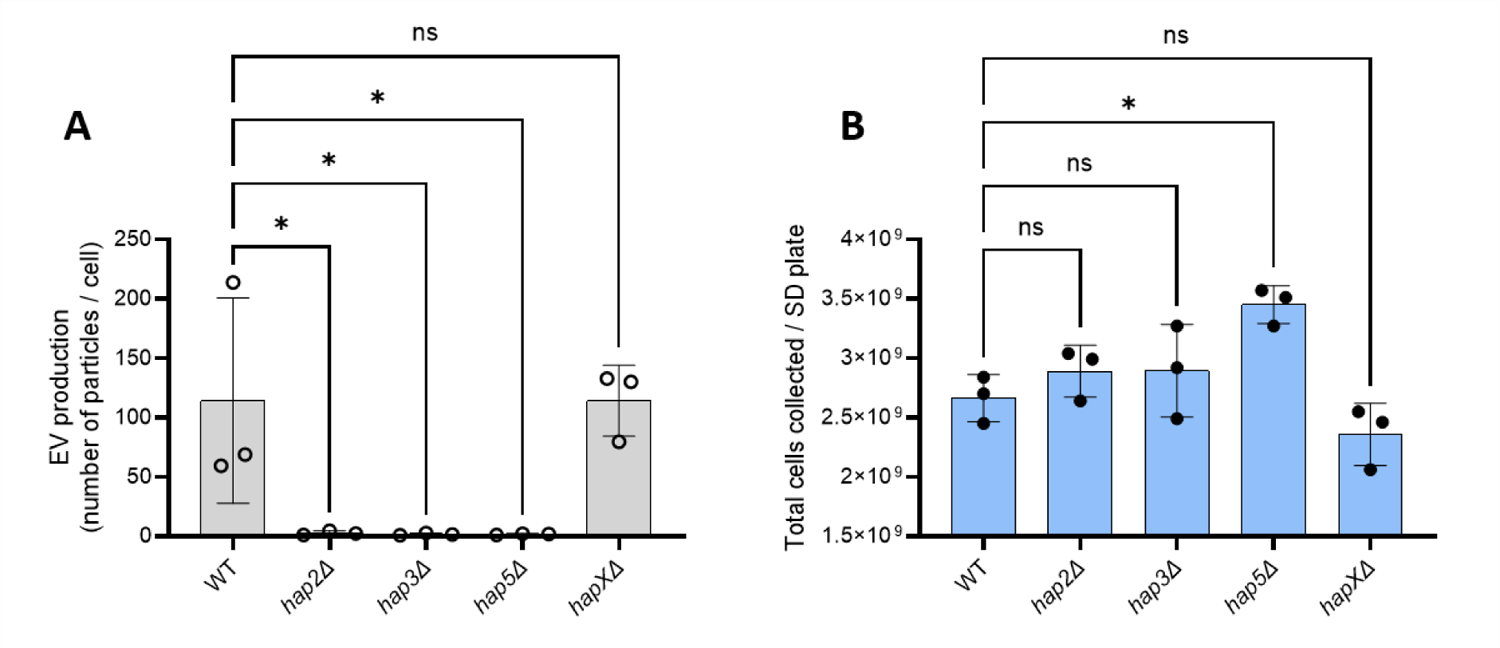
The *HAP2/3/5* complex is involved in EV secretion. Quantification of single EV particles released by WT and HAP complex mutants using nanoFCM (A). Growth analysis by counting the total number of cells obtained from SD plates (B). The experiments were carried out in biological triplicates. Error bars show means ± SD.

### c) Transcriptional analysis of EV-defective mutants

To decipher the link between the expression of these TF and the EV production phenotype of their corresponding mutants, we performed the RNA-seq analysis of the *hap2Δ*, *gat5Δ* and WT strains grown for 18 h on SD plates at 30°C, a condition which corresponds to the maximum EV production (see **Fig 1C**). Differential expression gene analysis identified 538 genes commonly upregulated in both *hap2Δ* and *gat5Δ* strains (**Fig 4A**) (**Supplementary Table S1**). Gene ontology (GO) analysis using the FungiDB database (https://fungidb.org/fungidb/app), revealed that genes associated with ribosome biogenesis, transmembrane transport, and protein-containing complex assembly were enriched in this set (**Fig 4B**). This suggests that the downregulation of the translation machinery expected to occur during the transition from the exponential phase to the stationary phase^29^ is impaired in both mutant strains. On the other hand, 310 genes were downregulated in both mutant strains (**Fig 4C**) (**Supplementary Table S1**). GO analysis revealed genes coding for proteins implicated in protein glycosylation, signaling, carbohydrate metabolic processes, cell wall organization and biogenesis, protein modification process, carbohydrate metabolism, reproductive processes, vesicle-mediated transport, as well as cytokinesis, as being enriched in this set (**Fig 4D**). Interestingly, *GAT5* expression is strongly downregulated (10-fold) in the *hap2Δ* mutant, suggesting that Hap2 might act upstream of Gat5 in regulating EV production. In contrast, neither *BZP2* nor *LIV4* appeared regulated by *HAP2* and/or *GAT5*, suggesting an independent regulation pathway. We then considered the top-100 most expressed genes in WT within the lists of *gat5Δ,* or *hap2Δ* downregulated genes reasoning that the impact of a gene downregulation associated with a mutation would have a phenotypic consequence if this gene is strongly expressed in the WT strain. Interestingly, 21 of the 39 previously identified *C. neoformans* enriched EV-protein encoding genes^14^ were regulated by either *GAT5* or *HAP2* (**Fig. 4E**). In addition, the 15 genes coding for EV proteins regulated by both TFs included seven proteins belonging to the top ten most abundant EV-associated proteins in *C. neoformans*. Thus, *HAP2* and *GAT5* regulate the expression of many EV-associated protein components, confirming their function as major regulators of EV production in *C. neoformans*.

**Figure 4:**
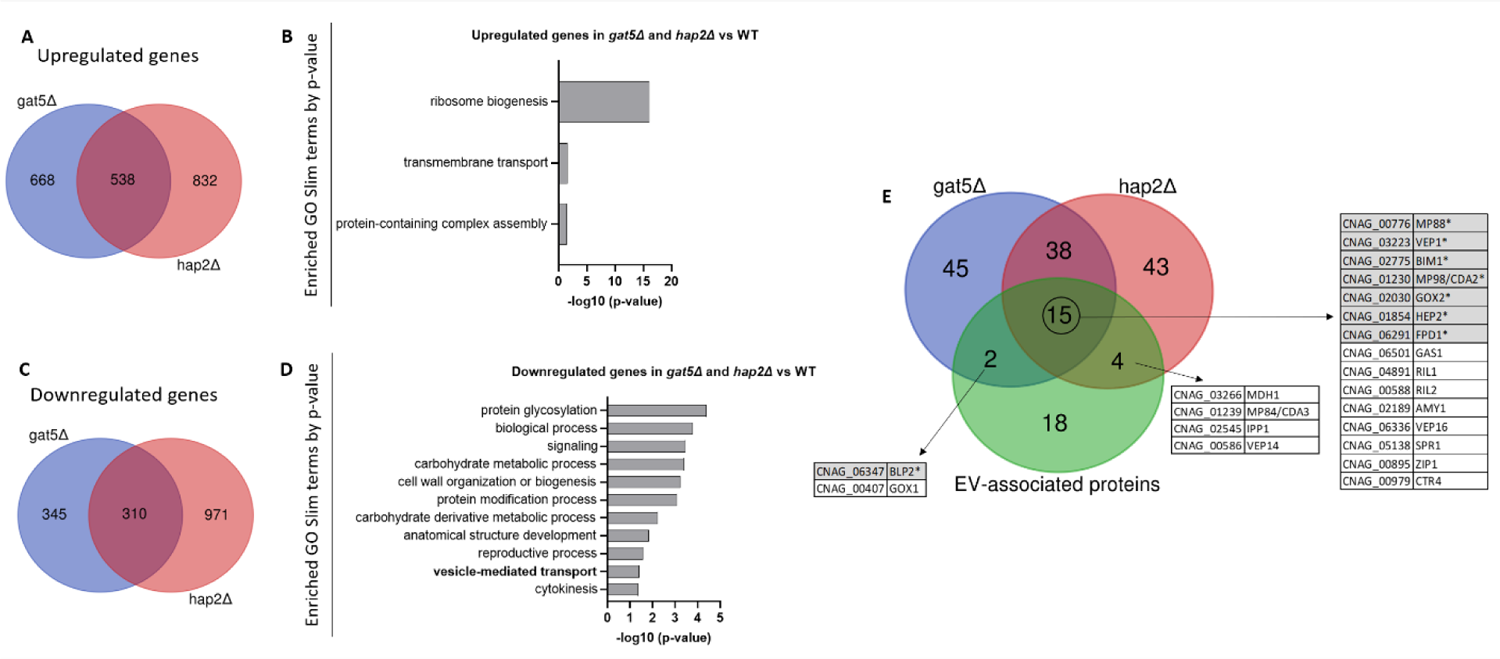
Transcriptomic analysis of the EV-deficient mutants *hap2Δ* and *gat5Δ* grown in EV-producing conditions. Venn diagram revealing the number of upregulated genes in both mutants (A). Analysis of the GO SLIM categories enriched in the group of genes upregulated in both mutants (B). Venn diagram revealing the number of the downregulated genes in both mutants (C). GO SLIM categories enriched in the group of downregulated genes in both mutants (D). Venn diagram revealing the overlap between genes downregulated in *gat5Δ* and *hap2Δ*, and the ones coding for the EV-enriched proteins of *C. neoformans*, previously identified by proteomics (E). The genes highlighted in grey with an asterisk encode proteins among the ten most abundant proteins in *C. neoformans* EVs.

### d) EV production regulator mutant strains are altered in azole susceptibility

Strikingly, all four newly identified EV production mutant strains have been reported to have altered FLC sensitivity in the systematic phenotypic analysis of the TF mutants published by the Bahn’s laboratory^25^. Indeed, we confirmed that *hap2Δ*, *gat5Δ,* and *liv4Δ* were less susceptible to FLC, while *bzp2Δ* was more susceptible compared to the WT strain (**Fig 5A-B**). Similar phenotypes were observed with Itraconazole and Voriconazole (**Fig 5A**). We also tested the FLC susceptibility of the *hap* mutants and noticed that those impaired in EV production (i.e., *hap2Δ, hap3Δ,* and *hap5Δ*) were also more resistant to FLC. In contrast, the *hapXΔ* mutant strain, which showed no alteration in EV production (**Fig3A**), only displayed slight differences in FLC sensitivity (**Fig 5B**).

**Figure 5:**
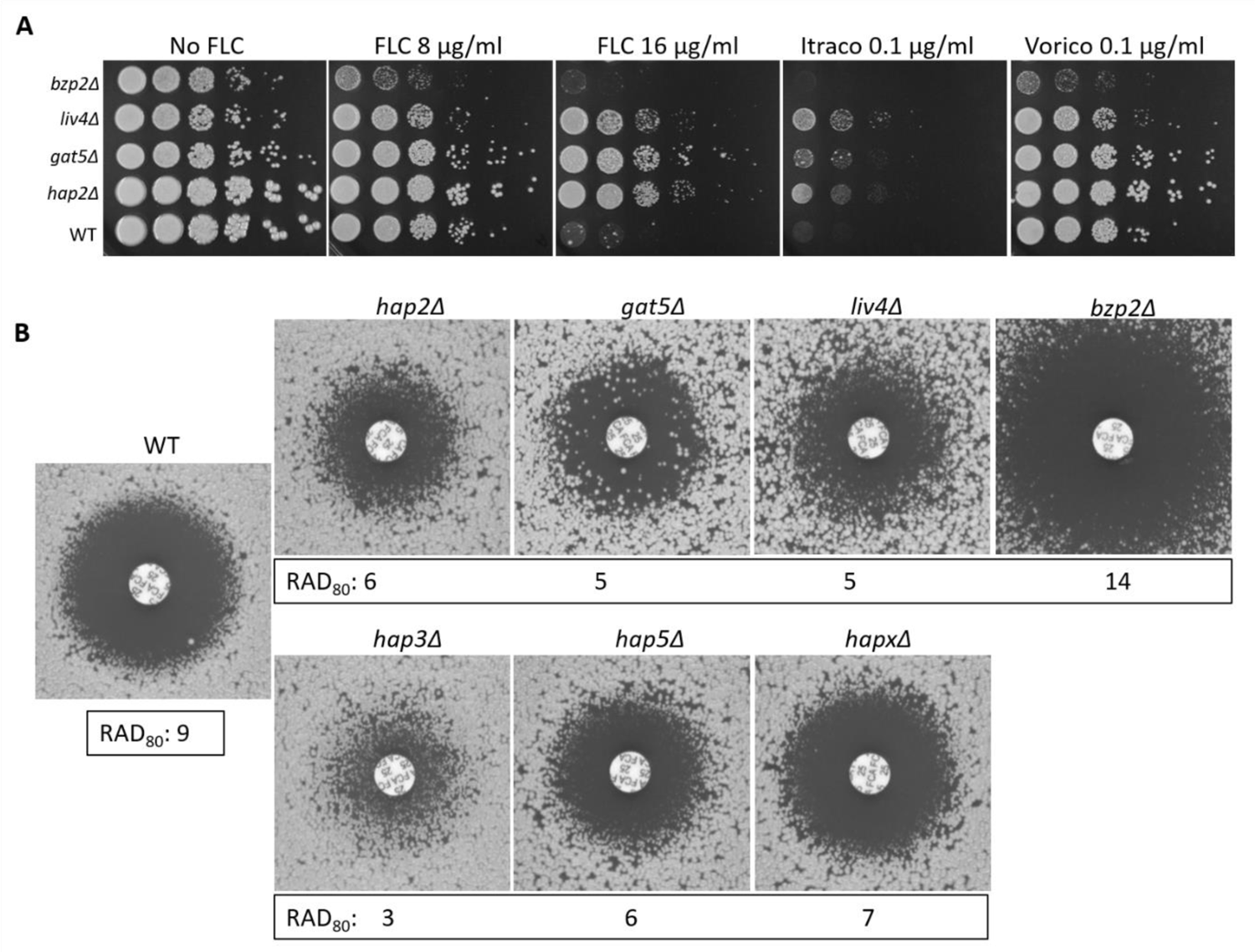
Transcription factors regulating EV release are associated with altered azole susceptibility. Spot assay for Fluconazole (FLC), Itraconazole (Itraco) and Voriconazole (Vorico) susceptibility (A) and FLC diffusion disk diffusion assay (25 µg/disk) for the four EV-defective TF mutants identified in the screening (*hap2Δ, gat5Δ, liv4Δ* and *bzp2Δ*), and for all the members of HAP complex inlcuing *hap3Δ*, *hap5Δ* and *hapXΔ,* grown for 72h (B). Average RAD_80_ values obtained from diskImageR analyses are provided. The results are illustrative of three biological replicates.

Accordingly, reconstruction of the *gat5Δ* and *hap2Δ* mutant strains restored both phenotypes to WT levels (**Fig 6A-D**). We also evaluated the sensitivity of these strains to agents that disturb the cell wall and plasma membrane, such as Calcofluor white - CFW, hydrogen peroxide - H_2_O_2,_ and sodium dodecyl sulfate – SDS, and observed that only *gat5Δ* was more susceptible to the presence of SDS (**Fig 6E**). Overall the analysis of TF mutant strains suggested that EV production and FLC susceptibility are coregulated.

**Figure 6:**
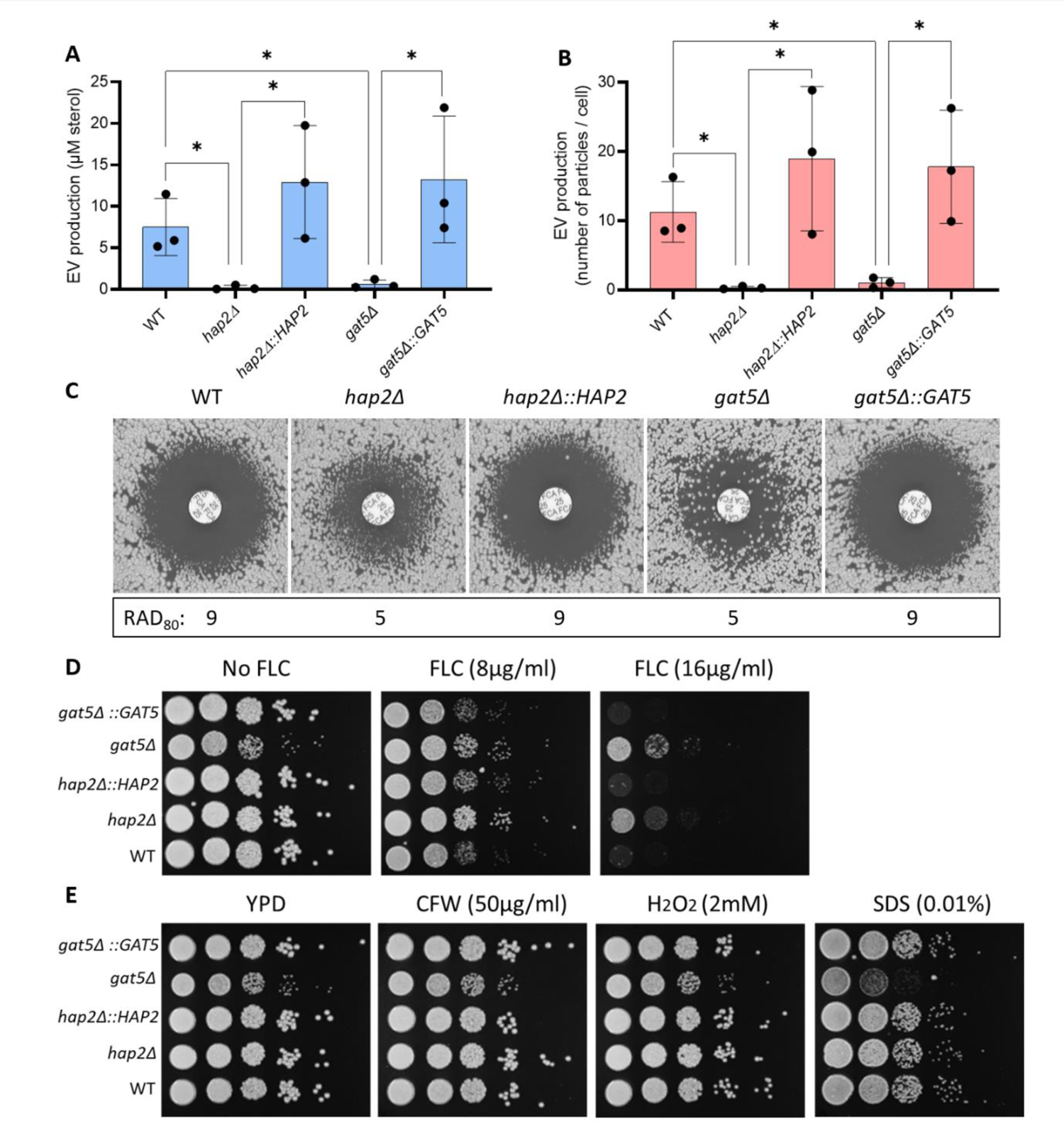
Complementation of the *hap2Δ* and *gat5Δ* mutant strains restores the phenotypes of EV production and FLC susceptibility. Total sterol contents in EVs produced by WT, mutants, and complemented cells. Sterol concentration values are expressed per 10^9^ cells for each condition (A). Quantification of EV particles using nanoFCM in WT, mutants and complemented strains (B). FLC susceptibility by disk diffusion assay (25µg/disk) and RAD_80_ values obtained by diskImageR (C). FLC susceptibility analysis by spot assay (D). Growth analysis in the presence of cell wall and plasma membrane disturbing agents (Calcofluor White - CFW, Hydrogen Peroxide - H_2_O_2,_ and Sodium Dodecyl Sulfate – SDS (E). The experiments were carried out in biological triplicates. Error bars show means ± SD.

### e) *hap2Δ* FLC resistance is not associated with *ERG11* or *AFR* genes over expression

In *C. neoformans,* FLC resistance has been associated with either mutation of *ERG11* (CNAG_00040) and/or the overexpression of efflux pumps genes such as *AFR1* (CNAG_00730)*, AFR2* (CNAG_00869) and *AFR3* (CNAG_06909)^30^. Interestingly, differential gene expression analysis of *gat5Δ, hap2Δ,* and WT cells grown under conditions of EV production revealed that neither mutation was associated with an increased expression of any of these resistance genes. We then used RT-qPCR assays to evaluate the expression levels of *ERG11* and the *AFR* genes in the presence or absence of FLC in these genetic backgrounds. As an additional control, we also included a strain lacking *NRG1* (CNAG_05222), which was shown to be FLC resistant^25^ but was found here to produce WT EV levels (**Fig S2**). In the absence of FLC, we did not observe changes in *ERG11* expression associated with either *HAP2* or *GAT5* deletion (**Fig 7A**). In the presence of the drug, the expression of *ERG11* in the *hap2Δ* mutant strain was similar to the WT, whereas in *gat5Δ*, we observed a two-fold reduction expression. In contrast, in the *nrg1Δ* mutant strain, we observed a two-fold overexpression of *ERG11* as compared to the WT strain. Thus, the FLC resistance phenotype in *hap2Δ* and *gat5Δ* mutants cannot be explained by an alteration in *ERG11* expression levels (**Fig 7A**).

**Figure 7:**
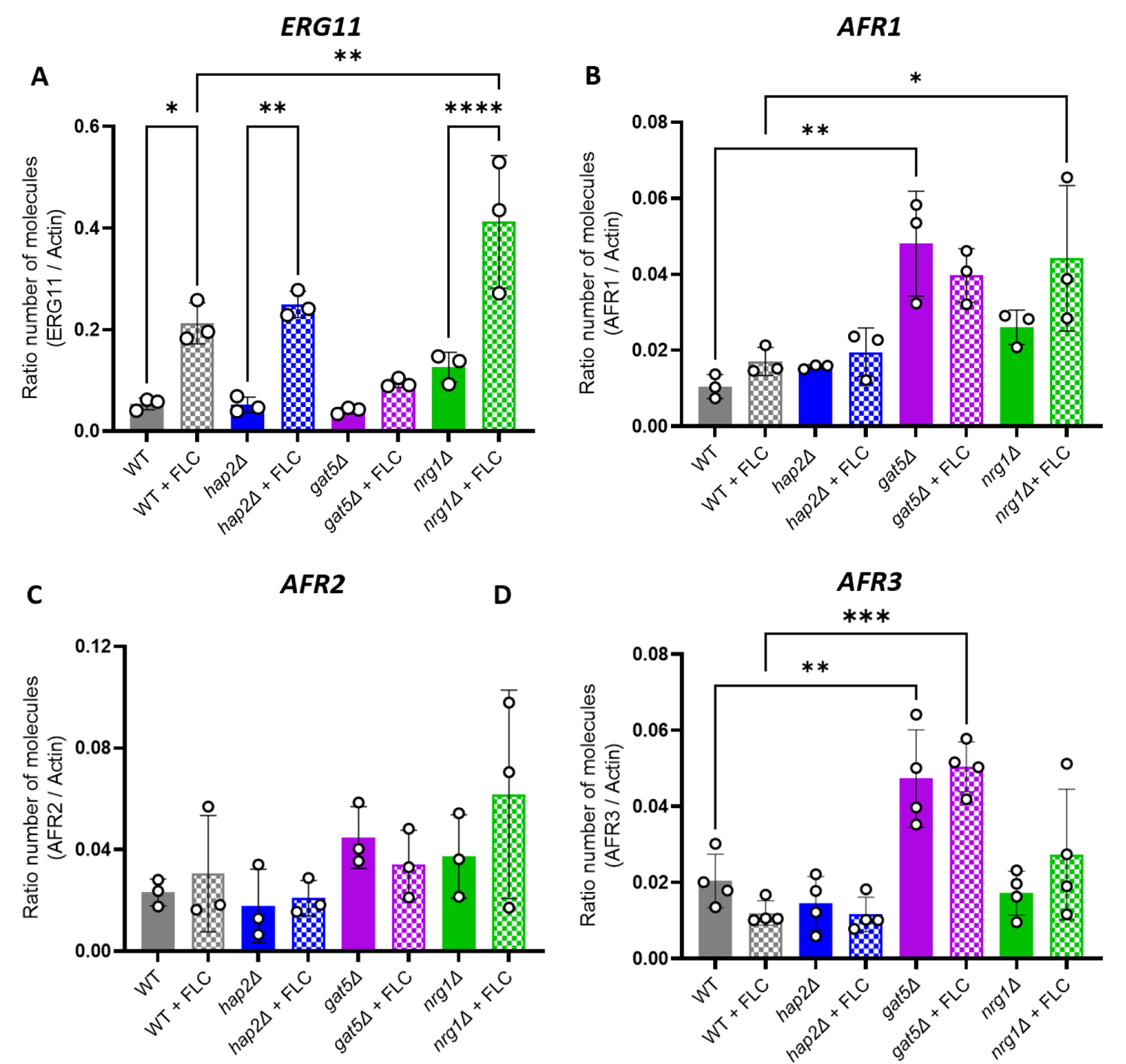
Gene expression analysis of known FLC resistance regulators in *C. neoformans*. *ERG11* (A), *AFR1* (B), *AFR2* (C), and *AFR3* (D) gene expression profile as generated by RT-qPCR. The experiments were carried out in at least biological triplicates. Error bars show means ± SD.

In contrast to *ERG11*, the presence of FLC did not impact the expression of *AFR* genes in any of these strains (**Fig 7B–C–D**). However, *AFR1* and *AFR3* but not *AFR2* were upregulated in the *gat5Δ* mutant (both with and without FLC), suggesting that these two efflux pumps could play a role in driving FLC resistance in this strain (**Fig 7B–C–D**). A similar result was obtained with the *nrg1Δ* mutant, although to a lesser extent for *AFR3*. Interestingly, in the *hap2Δ* strain, the expression of the *AFR* genes wasnot altered, suggesting that additional mechanisms are driving FLC resistance in these cells.

### f) FLC regulates EV production and cellular lipid profile

The analysis of the EV regulators suggests that changes in EV production impact FLC resistance. We then studied the converse relationship and tested whether FLC could modulate EV production in *C. neoformans*. We first tested the growth of WT cells on SD containing FLC at different concentrations (0.3, 0.6, 1.25, 2.5, and 5 µg/mL) identifying 0.6 µg/mL as the highest FLC concentration, which was not affecting cellular growth (**Fig 8A**). At this concentration, the expression of *ERG11* was not altered (**Fig 8B**). However, EV production was reduced by 2.4-fold (**Fig 8C**), suggesting that FLC regulates EV production levels.

**Figure 8:**
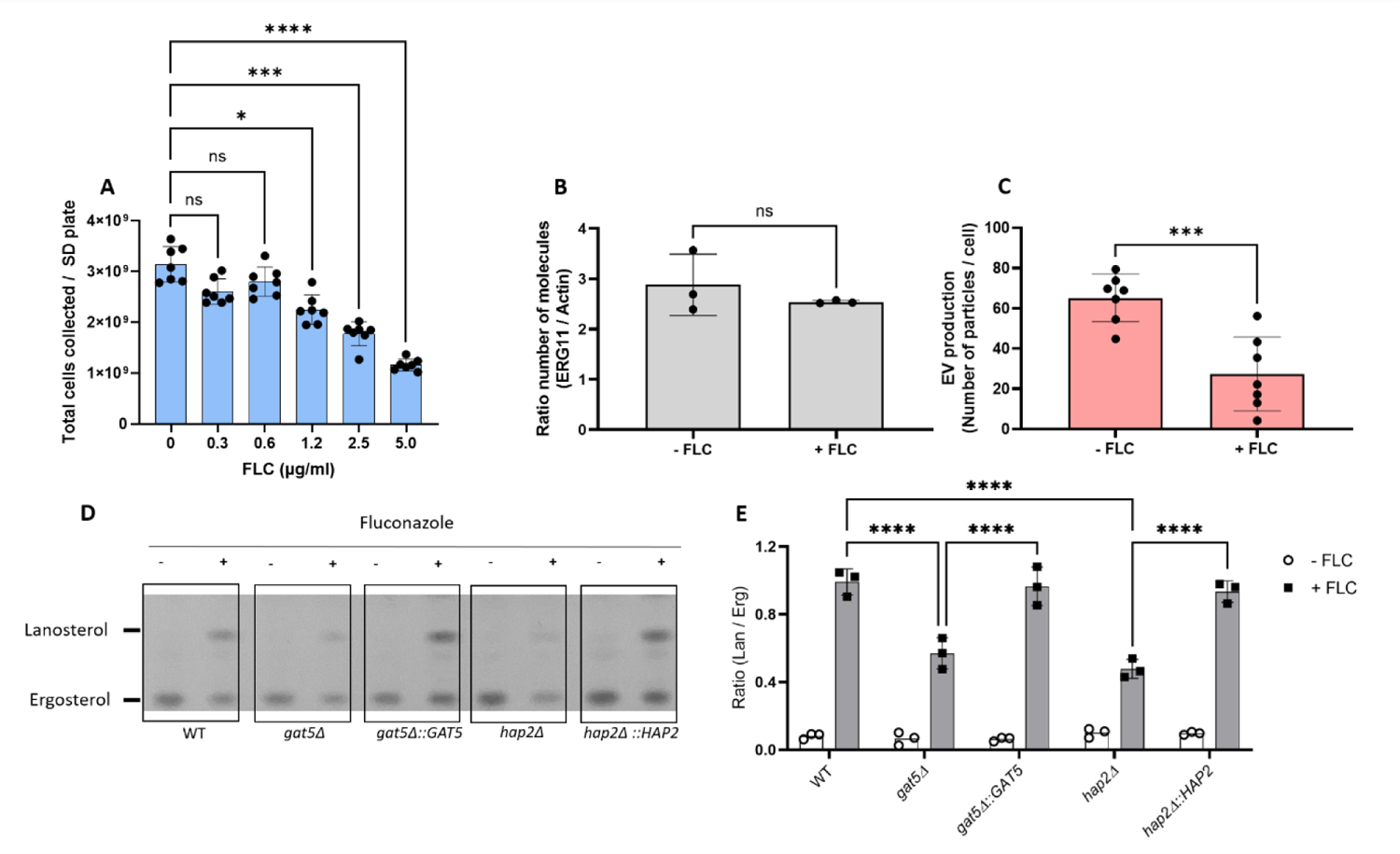
Fluconazole treatment affects EV production and cellular sterol homeostasis. WT cells were grown in different concentrations of FLC on SD agar medium, and the total number of cells was counted (A). Levels of *ERG11* expression in the presence of a subinhibitory concentration of FLC (0.6 µg/mL) (B). Analysis of EV production by nanoFCM in the presence of FLC (0.6 µg/mL) (C). Analysis of lanosterol and ergosterol lipid profile by thin layer chromatography, TLC in cells (D). Analysis of TLC bands by densitometry units and ratio between lanosterol and ergosterol-specific signals for cells (E). The experiments were carried out in at least two biological duplicates, and representative results were shown. Error bars show means ± SD.

FLC is an inhibitor of the Erg11 lanosterol 14-demethylase, which transforms lanosterol in 4,4-dimethyl-cholesta-8,14,24-trienol in the fungal ergosterol biosynthetic pathway^31^. We thus compared the cellular lipid profile in WT and EV-defective mutant cells in the presence or absence of the same FLC-subinhibitory concentration (0.6 µg/mL). As expected^32^, FLC treatment led to lanosterol accumulation in all strains under the growth conditions examined (**Fig 8D)**. However, decreased accumulation was observed in the *gat5Δ* and *hap2Δ* mutant strains as revealed by the ratio of lanosterol/ergosterol signals, suggesting that the lipid composition is altered in these EV-defective strains (**Fig 8E**).

### g) FLC-resistant isolates obtained *in vitro* are impaired in EV production

To further explore the link between FLC resistance and EV production in *C. neoformans*, we analysed the EV production in spontaneous FLC-resistant isolates obtained *in vitro*. Previous studies reported that when *C. neoformans* cells are spread on a medium containing an inhibitory FLC concentration, a subset of the cell population can grow and produces colonies. This phenomenon is known as heteroresistance and was associated with the duplication of chromosome 1 bearing *ERG11* and *AFR1* resistance genes^12, 31, 33^. This FLC-resistant phenotype can be reverted by subculturing the cells in a drug-free medium, the phenotypic reversion being associated with the reversion of the aneuploid genotype^11^. Therefore, we isolated independent spontaneous FLC-resistant colonies on plates containing 15 µg/mL of FLC. FLC-susceptible parental and FLC-resistant isolates were then passaged six times in a liquid drug-free YPD. A schematic overview of the experimental design is delineated in **Fig 9A.** All parental, resistant, and passage strains were tested for EV production and FLC sensitivity. As expected, the sixteen passage 0 (P0) strains isolated on plates containing FLC were resistant to FLC, as revealed by a significant decrease in the inhibition halo in FLC-disk diffusion assays (**Fig 9B** and **9D**). Interestingly, this drug-resistant phenotype was accompanied by a substantial reduction in EV production as analysed by nano-FCM (**Fig 9C**). Importantly, none of the sixteen FLC-resistant isolates had pronounced growth defects in the SD medium, presenting the same transition time frame from the exponential to stationary phase (**Fig S3**). In contrast, the passage 6 (P6) strains all lost their resistant phenotype, although to a different extent (**Fig 9E** and **9G**), and they also restored EV production to parental levels (**Fig 9F**).

**Figure 9:**
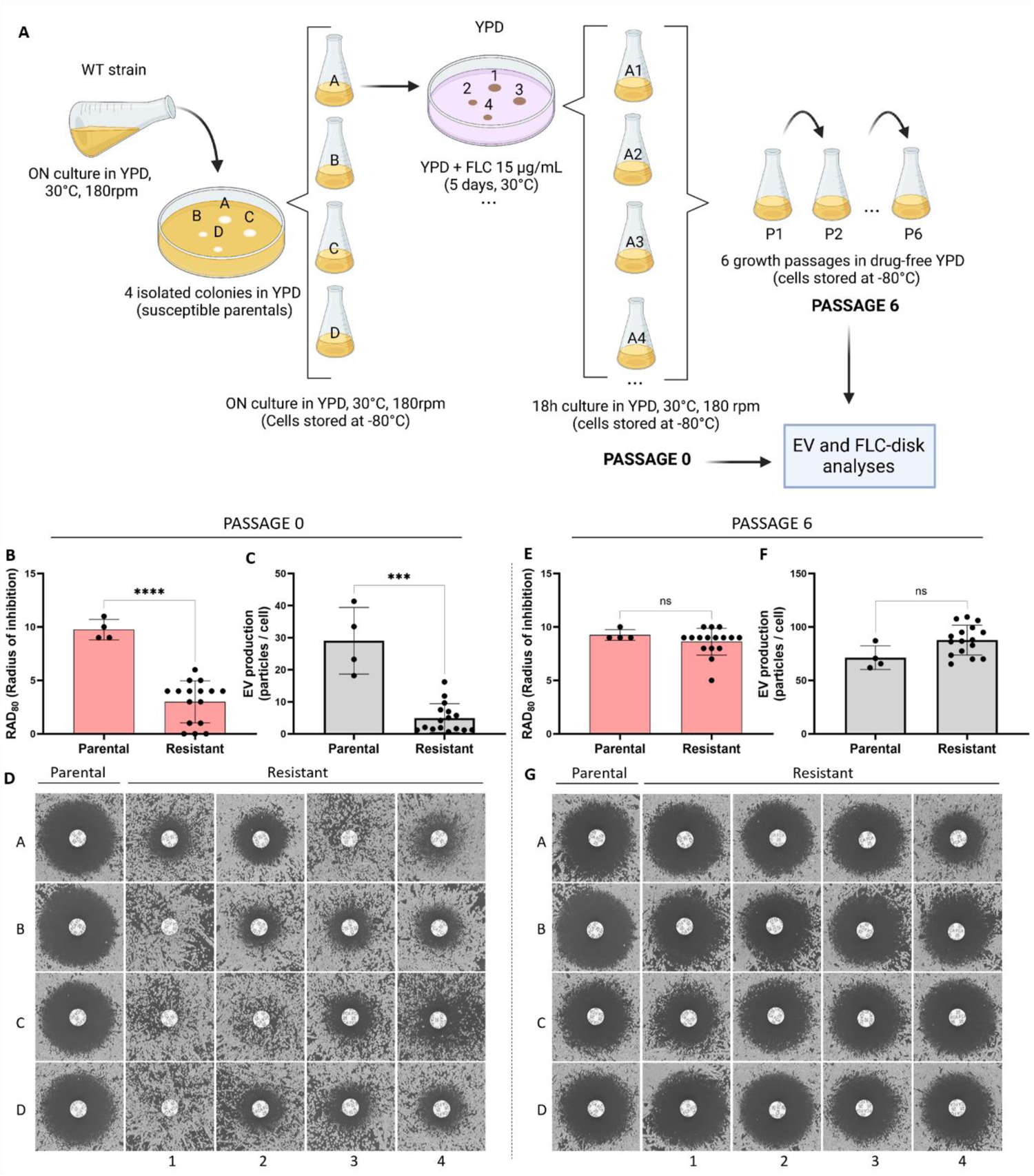
Analysis of FLC susceptibility and EV production in FLC-resistant isolates obtained *in vitro*. Illustrative scheme of the experimental strategy used to explore the association between FLC resistance and EV production in spontaneous drug-resistant strains obtained *in vitro* (A). Cells at passage 0: quantitative analysis of the growth inhibition halo from the FLC-disk diffusion assays by diskImageR, RAD_80_ (B), EV production analysis by nanoFCM (C), and qualitative analysis of the inhibition halo of the FLC-disk diffusion assays (D). Cells after six successive passages in the absence of FLC: quantitative analysis of growth inhibition halo from the FLC-disk diffusion assays by diskImageR, RAD_80_ (E), EV production analysis by nanoFCM (F), and qualitative analysis of the growth inhibition halo (G). Schematic representation created in BioRender.

We then evaluated the chromosome copy numbers of the parentals, FLC-resistant, and passaged strains by qPCR (using primers for genes located on chromosome 1 left arm (CNAG_00047), chromosome 1 right arm (CNAG_00483) and chromosome 2 (CNAG_03602)). As expected, all parentals were monosomic for chromosome 1 (**Fig 10A and 10D**). In agreement with the published literature, we also found that fourteen out of sixteen FLC-resistant strains were disomic for chromosome 1 (**Fig 10B** and **10E**). Meanwhile, all P6 strains reverted to the original monosomic state of chromosome 1 (**Fig 10C** and **10F**).

**Figure 10:**
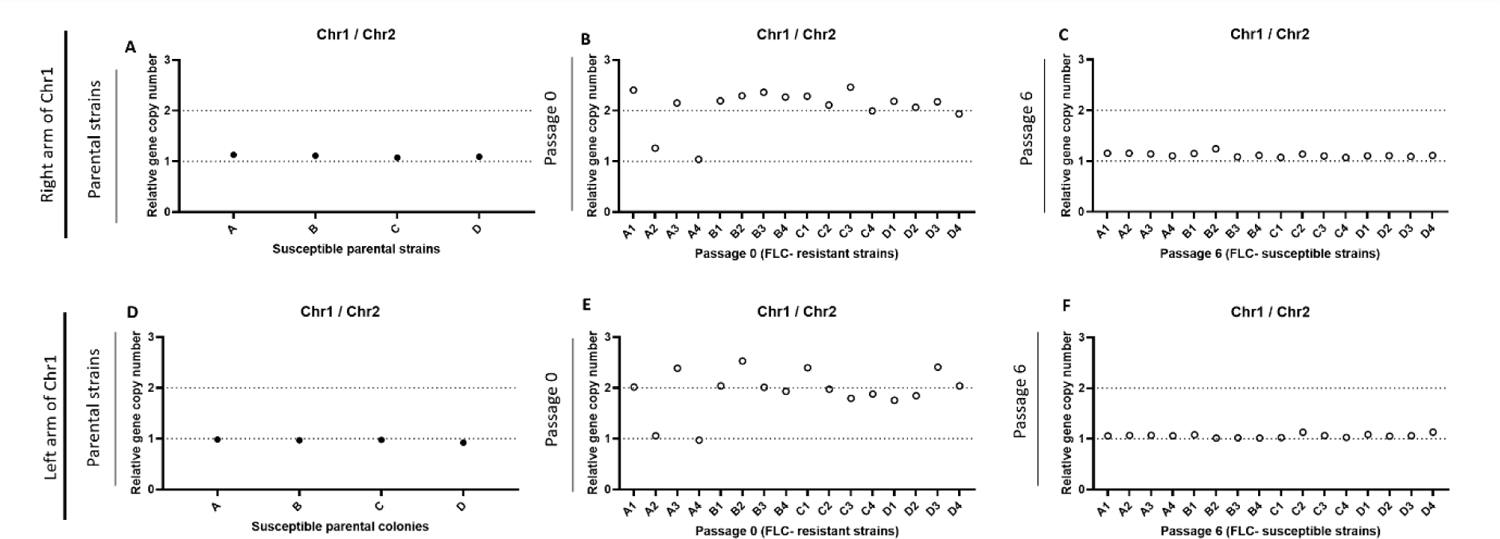
qPCR assays evaluating the monosomic or disomic status of chromosome 1 in the FLC sensitive and resistant strains obtained *in vitro*. Parental strains (A and D), FLC-resistant passage 0 strains (B and E), and passage 6 strains (C and F) were tested using primers specific for genes located on chromosome 1 right (A to C) and left arms (D to F). Primers specific to chromosome 2 were used as control.

We sequenced the genomes of the A series to examine possible copy number variations. This analysis confirmed that strains A1 and A3 were disomic for chromosome 1, while strains A2 and A3 were disomic for chromosome 4 (**Fig 11A**). Interestingly, while *ERG11* but not *AFR1* was overexpressed in the P0_A2 isolate, we found no change in the expression of these resistance genes in the P0_A4 isolate, suggesting that its FLC-resistant phenotype was independent of their regulation (**Fig 11B**-**C**).

**Figure 11:**
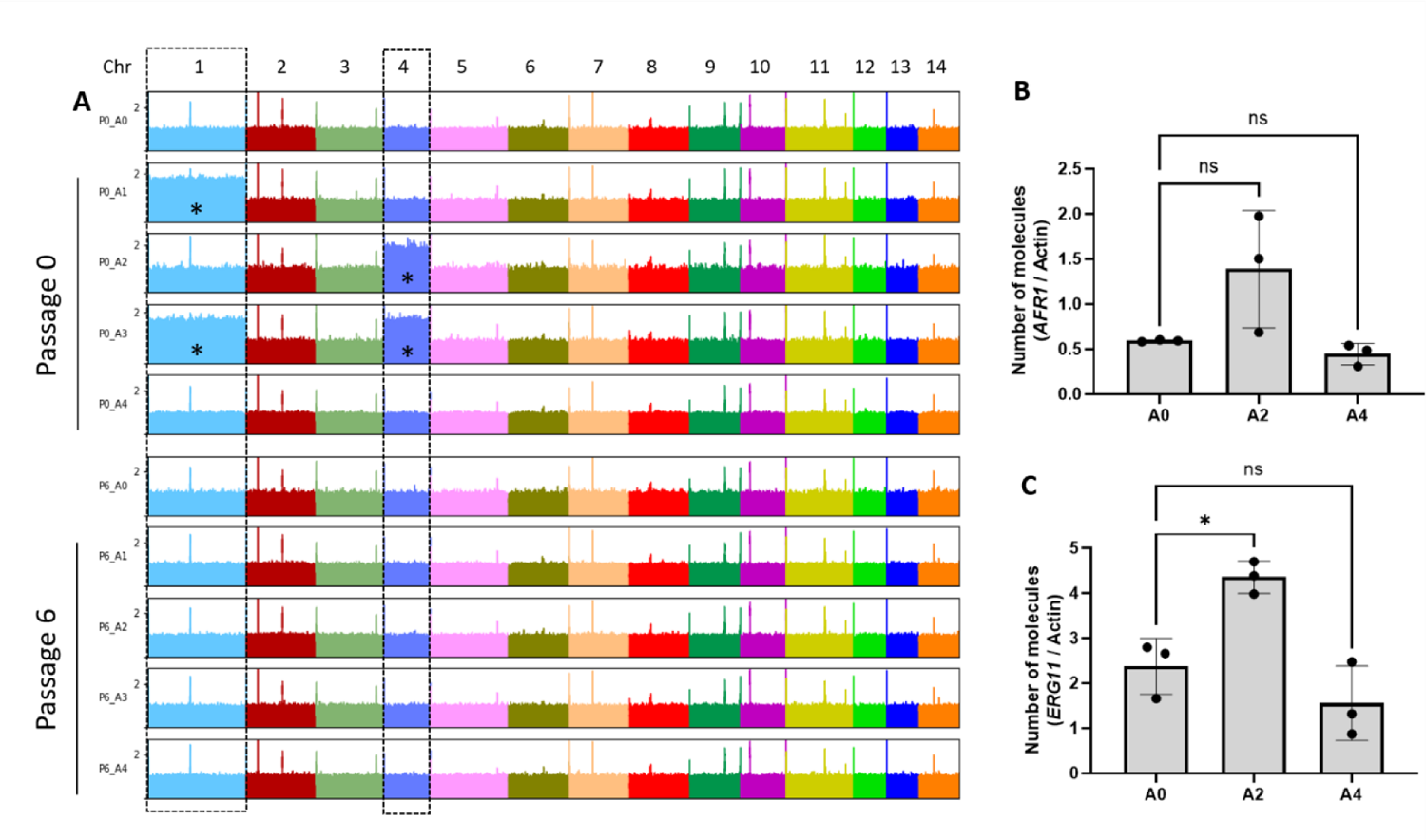
Chromosome duplications revealed after aligning DNA sequencing reads obtained from the A series of strains to the *C. neoformans* reference genome. (A). Analysis of *AFR1* (B) and *ERG11* (C) gene expression in A0, A2, and A4 strains by RT-qPCR. Indicative of chromosome duplication (*).

### h) FLC resistance and regulation of EV release in *C. neoformans* clinical isolates

To test whether the relationship between EV production and FLC resistance would extend to clinical isolates, we studied a series of isolates having different levels of FLC susceptibility. Plotting the the radius of inhibition (RAD_80_) from FLC-disk diffusion assays and the average levels of EV production from these isolates did not reveal a clear correlation (R^2^ = 0.002), as both the fully resistant isolates (no inhibition halo), as well as the most sensitive ones, were poor EV producers (**Fig 12A**). We then considered serial isolates of *C. neoformans* that had become resistant during infection. For the isolates CNRMA17.247 and CNRMA20.738, we observed that the acquisition of FLC resistance coincided with the decreased ability to produce EVs (**Fig 12B**). For the second lineage (isolates CNRMA4.1291 and CNRMA4.1293), we did not observe any difference in EV production between the sensitive and the resistant isolates (**Fig 12C**). Interestingly, SNPs analysis revealed a CNRMA4.1293 specific (A to T) nonsynonymous mutation in *ERG11*, responsible for a Y145F mutation and possibly associated with the acquired resistance^34^. We then tried to revert the resistance phenotype of four FLC-resistant clinical isolates (CNRMA20.738, CNRMA4.1293, CNRMA4.158, and CNRMA5.114) by growing them in a drug-free YPD medium for eight subsequent passages (P8). CNRMA5.114 was the only isolate for which we obtained a more sensitive derivative (**Fig 12D**). In agreement with our previous observation, the P8-sensitive passage produced more EVs than the original P0-resistant isolate (**Fig 12E**). As expected, P8 strains with no change in drug susceptibility also did not display altered EV production (**Fig 12D-E**). Sequencing the genomes of the two lineages of recurrent isolates as well as the P0 clinical isolates and their P8 derivatives revealed no chromosomal duplications (**Fig S4**). Among the sequenced P0 clinical isolates, we observed the disomy of chromosome 12 in CNRMA5.114, which was lost in its P8 sensitive derivative. Partial duplication of chromosome 14 was also observed in the reverted P8 strain (**Fig 12F**). We did not further investigate a possible causative link between FLC resistance and the observed genotype modifications in these isolates.

**Figure 12:**
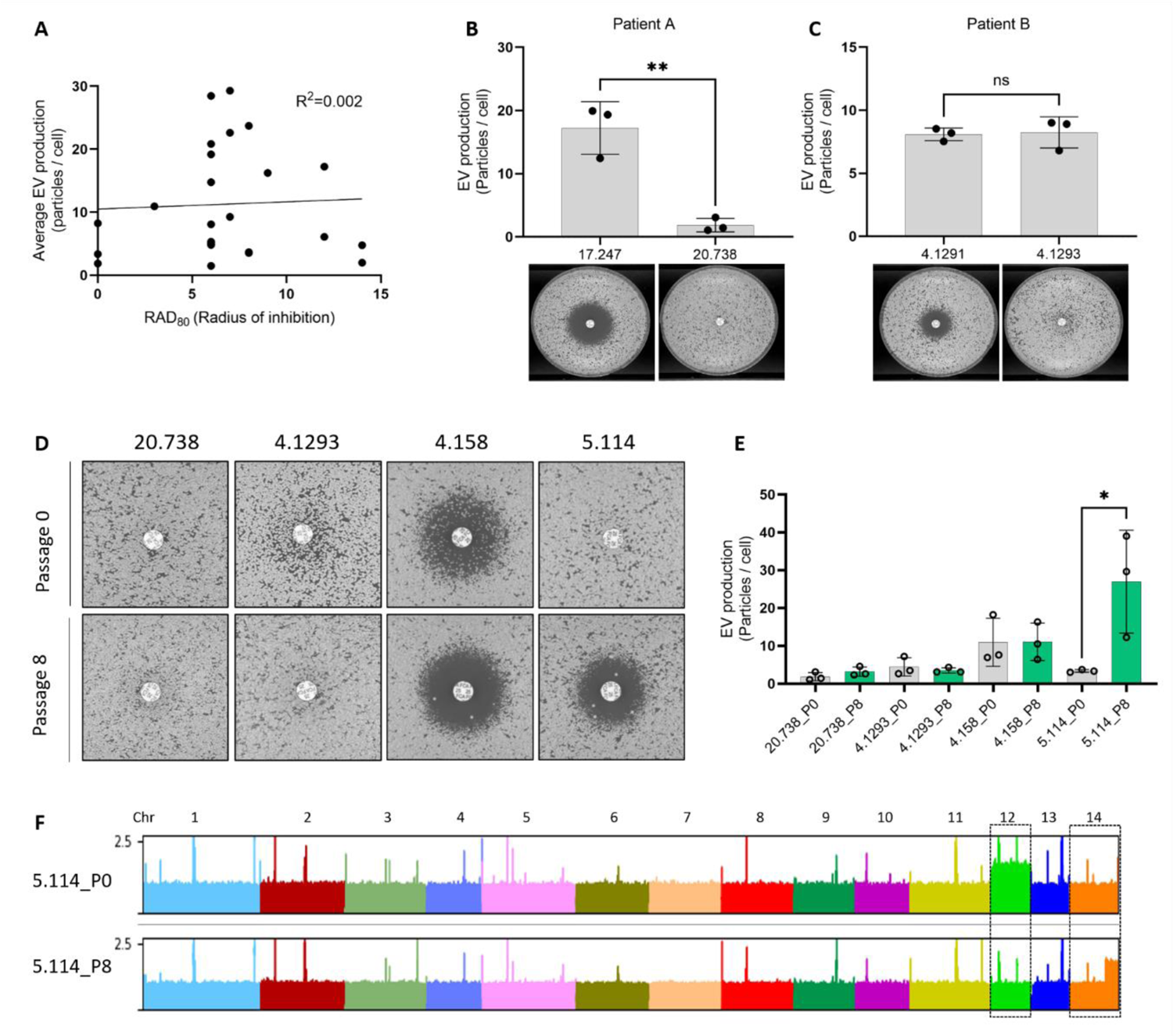
EV production and FLC resistance are linked in clinical isolates. Analysis of EV production by nanoFCM and size of FLC-disk inhibition halo in clinical isolates by linear regression (A, radius of inhibition = RAD_80_ values by diskImageR). EV production as measured by nanoFCM in the recurrent isolates CNRMA17.247 and CNRMA20.738 (top) and FLC-disk diffusion assays (bottom) from patient A (B). EV production as measured by nanoFCM in the recurrent isolates CNRMA4.1291 and CNRMA4.1293 (top) and FLC-disk diffusion assays (bottom) from patient B (C). FLC-disk diffusion assay in passaged clinical isolates (D) and analysis of EV production of passaged clinical isolates by nanoFCM (E). Alignment of DNA-Seq reads obtained from P0, and P8 CNRMA5.114 passaged clinical isolates (F). EV and FLC-disk analyses were carried out in at least biological triplicates, and representative results were shown.

## 3) Discussion

The mechanisms of biogenesis of EVs in fungi are still unknown. In this paper, we present experiments revealing a tight regulation of EV production in response to changes in growth conditions in *C. neoformans,* confirming previous reports in other models^35, 36^. We also characterize four transcription factor mutant strains strongly impaired in EV production which are the first to be described in fungi and, to our knowledge, in any organisms. Surprisingly, we observed no major growth defect associated with *hap2Δ* mutation, suggesting that a WT level of EV production is dispensable in this fungus, at least under the examined conditions.

Our experiments also indicate that EV production is also strongly regulated by fungal growth phase. EVs are produced between 16 h and 22 h after cell plating, corresponding to the transition between the exponential phase and stationary phase. This result was consistent with reports of cells growing on YPD solid medium^36^. In *S. cerevisiae*, the transition from the exponential phase to the stationary phase is associated with a shift from fermentation to respiration and a sharp slowdown of the general metabolism^37, 38^. The Hap complex is the major regulator of this shift, controlling respiration and mitochondrial function in this yeast^39, 40^. In pathogenic fungi, this complex has been studied mostly for its regulatory function in iron metabolism^41–43^. As in *S. cerevisiae* ^44^, the Hap complex seems to control respiration in *C. neoformans*. Thus, Hap3 and HapX have also been shown to negatively regulate genes encoding respiratory and TCA cycle function under low iron condition^45^. The *HAP* genes have also been shown to regulate sexual development in association with the pheromone-responsive Cpk1 mitogen-activated protein kinase pathway in this pathogenic yeast^40^. However, phenotypic analyses of the *hap* mutants suggest that HapX and the Hap2/3/5 complex might also have independent functions. For instance, *HAP3* and *HAP5* but not *HAPX* are necessary for growth on ethanol^43^, and the Hap2/3/5 complex, but not HapX, is crucial in repressing pheromone production and cell fusion during mating^40^. Similarly, we found that the Hap2/3/5 complex, but not HapX, controls FLC resistance and EV production in *C. neoformans*.

The FLC resistance phenotype of the *hap* mutant strains is consistent with the literature^45, 62^. Counterintuitively, Hap3 and HapX have been reported to positively regulate several ergosterol biosynthetic genes^45^. Our differential gene expression analysis, which used different growth conditions, identified only two genes, *ERG20* (CNAG_02084) and *ERG27* (CNAG_07437) downregulated and one upregulated, *IDI1* (CNAG_00265) upon *HAP2* deletion. Overall, these data suggest that Hap2 in *C. neoformans* regulates global cellular changes impacting fluconazole resistance beyond the regulation of the ergosterol pathway.

In this work, we present cumulative lines of evidence suggesting a causative link between FLC resistance and EV production. First, all four EV production mutants displayed altered FLC susceptibility. We also observed this association with other hap mutants; *hapxΔ* being the only one producing WT level of EVs yet displaying a slight change in FLC suceptibilty. Second, all spontaneous FLC-resistant mutant strains isolated *in vitro* reduced the EV production, and both phenotypes reverted jointly. As many spontaneous FLC mutant strains are disomic for chromosome 1 (known to bear *ERG11* and *AFR1*^12^), one can imagine that repressors of EV production could also be located on this chromosome. Thus, chromosome 1 disomy could increase the expression of both FLC-resistant genes and EV repressors. However, we also identified two FLC-resistant spontaneous mutant strains with no duplication of chromosome 1; one being disomic of chromosome 4. Interestingly, none of the EV production regulators identified here are located on chromosome 1.

Finally, although we did not observe any correlations between the level of drug resistance and EV production among clinical isolates, the analysis of a couple of serial isolates from the same patient for which no mutation classically involved in FLC resistance could be identified revealed that the resistant strain produced lower EV amounts compared to the sensitive one. Notably, we did not observe any changes in chromosome copy numbers in these strains. Moreover, a sensitive strain derived from a resistant clinical isolate obtained after several passages in a drug-free medium produces more EVs than the original strain. This reinforces the idea that EV production and FLC resistance are strongly linked, at least in a subset of genetic backgrounds.

Nevertheless, the molecular mechanism by which Hap2 and Gat5 regulate both EV production and FLC susceptibility remains unknown. Overexpression of the drug efflux pumps *AFR1* and *AFR3* in *gat5Δ* could contribute to the resistance phenotype. Similarly, disruption of the lanosterol/ergosterol equilibrium in the presence of FLC in the EV-deficient mutants could also impact drug resistance. Indeed, a recent report revealed that chemical disruption of the plasma homeostasis is sufficient to modify azole resistance in pathogenic fungi^47^. However, it is unclear whether the decreased EV production alters plasma membrane homeostasis or whether these two TFs regulate membrane homeostasis and, consequently, EV production. Previous data in *Cryptococcus* suggest that the deletion of genes implicated in plasma membrane homeostasis can impact EV production and/or cargo loading^48, 49^. On the other hand, the fact that most EV structural protein-encoding genes are strongly downregulated in both *gat5Δ* and *hap2Δ* mutants suggests a direct effect of these TF on EV biosynthesis. The fact that a Y145F *ERG11* mutation regulates FLC resistance without altering EV production also favors this second hypothesis.

Interestingly, we found that a low concentration of FLC reduces EV production. Yet, this concentration was not sufficient to alter the *ERG11* gene expression, suggesting that cells might regulate EV production to maintain plasma membrane homeostasis. The impact of low antifungal concentrations on fungal cell biology is poorly studied ^6, 50^, but our data clearly indicates that fungal cells might regulate EV production as a first line of defense under membrane stress. In this model (**Fig 13**), FLC exposure triggers distinct types of cellular responses; *ERG11* overexpression and EV production would act to regulate plasma membrane homeostasis, whereas regulation of efflux pumps would minimize FLC concentrations in the cell.

**Figure 13:**
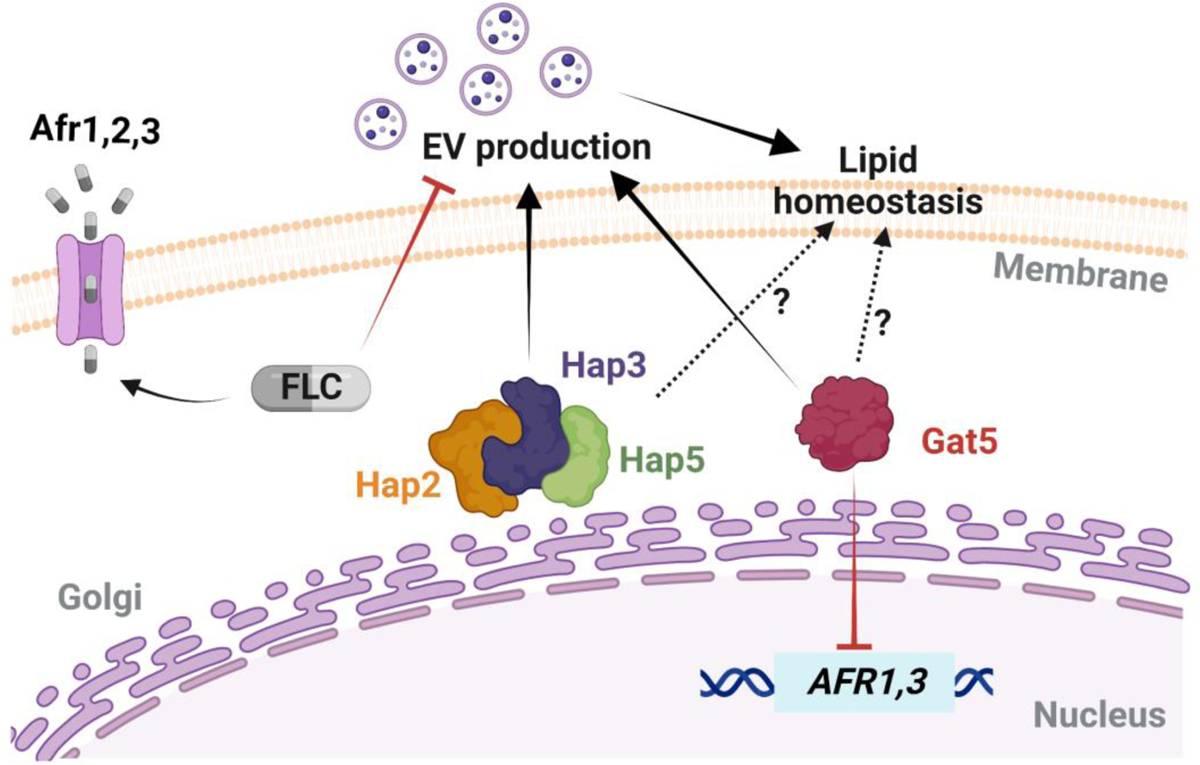
Model of EV biogenesis regulation and FLC susceptibility in *C. neoformans*. Specific gene mutations and fluconazole (FLC) exposure can trigger diverse cellular responses, including changes in the expressions of drug efflux-associated genes, of the azole target gene, and in extracellular vesicles (EV) biogenesis. Gat5 inhibits the expression of *AFR1* and *AFR3*, which can affect the intracellular FLC levels, while sub-inhibitory concentration of FLC can inhibit EV production. Hap2/3/5 and Gat5 regulate EV production, and cellular lipid homeostasis. The differential regulation of these cellular processes may determine FLC resistance phenotypes in this fungal pathogen. Schematic model created in BioRender.

Overall, this study identified the first TF regulating fungal EV biosynthetic pathways, while also uncovering novel roles that EVs play in modulating azole susceptibility. The genetics of EV production is still in its infancy, but functional genetics will further our understanding of EV biogenesis, their impact on drug resistance, and their roles during fungal infection.

## 4) Methods

### Media and culture conditions

All strains were taken from the G. Janbon laboratory collection at −80°C, plated onto yeast peptone dextrose (YPD) agar plates (1% yeast extract, 2% bacto-peptone, 2% dextrose, 2% bacto-agar) and incubated at 30°C for 48h before each experiment. Cell inoculum was performed in broth YPD (1% yeast extract, 2% bacto-peptone, 2% dextrose) rotating at 180 rpm for 24h at 30°C. The other solid media used in this study included Sabouraud (SAB) containing 1% bacto-peptone, 4% dextrose, 2% bacto-agar, Synthetic dextrose (SD) containing 0.67% yeast nitrogen base (Difco) without amino acids, 2% glucose and 2% bacto-agar; and Capsule induction medium (IM) containing 1.7 g of yeast nitrogen base without amino acids and without ammonium sulfate (Difco), 1.5 g of asparagine and 20 g of glucose L^-1^ of buffer (12mM in NaHCO_3_, 35mM in MOPS, pH 7.1), as previously described^51^. Drug-containing plates were made by adding a stock solution of 3 mg/mL FLC powder in water to different final concentrations depending on each experiment. The drug was added to the agar after autoclaving, then cooled to approximately 50°C before pouring.

### Fungal strains and mutant construction

The *C. neoformans* TF mutant library^26^ was obtained from the Fungal Genetic Stock Center (USA). This collection has been constructed in an H99o background^52^. After the screening, the phenotype of the identified EV mutant strains was thus confirmed in the KN99α^53^ background. All the following experiments have been performed in KN99α reference background^52^. The strains used are listed in **Supplementary Table S2**. The strains *MATα hap2Δ::NAT* (CNAG_07435), *MATα hap3Δ::NAT* (CNAG_02215), *MATα hap5Δ::NAT* (CNAG_07680), *MATα hapXΔ::NAT* (CNAG_01242), *MATα gat5Δ::NAT* (CNAG_05153), *MATα nrg1Δ::NAT* (CNAG_05222) have been constructed in the Hiten Madhani lab (UCSF, USA) and obtained from the Fungal Genetic Stock Center. The genotypes of these mutant strains were confirmed by PCR using gene-specific internal primers. To construct the strains NE1539 (*MATα bzp2Δ::NAT*) and NE1540 (*MATα liv4Δ::NAT*), we replaced the entire CNAG_04263 (*BZP2*) and CNAG_06283 (*LIV4*) CDS by the NAT marker. The deletion cassettes construction and transformant screening were performed as previously described^33^. The *GAT5* and *HAP2* genes were re-introduced into the corresponding mutant strain using a transient CRISPR-Cas9 expression (TRACE) system^54^. DNA fragments spanning a genomic region from 1kb upstream to 1kb downstream the CDS of the genes were PCR amplified and cloned into the pSDMA57 plasmid before being integrated at the genomic “safe haven” locus in *gat5Δ* and *hap2Δ* mutants, repectively^55^. All plasmids and primer sequences used are provided in **Supplementary Tables S3 and S4**, respectively. Twenty-four clinical isolates of *C. neoformans* were also used. All isolates were recovered from the national cryptococcosis surveillance program managed by the National Reference Center for Invasive Mycoses & Antifungals (NRCMA, Institut Pasteur) between 2004 and 2021. All the clinical isolates are provided in **Supplementary Table S2.**

### EV isolation protocol and optimization for EV screening in 96-well plates

EV isolation by the conventional ultracentrifugation method followed the steps of the previously described protocol^14^. A 96-format protocol was also used to test a large number of samples concomitantly. For this, cells from the stock were steacked on YPD agar plates for 48h at 30°C, and one loop of cells was inoculated in 1 ml of YPD broth of each well of a 96-deep well plate and incubated for 18h at 30°C under agitation. The plate was centrifuged, and cells were washed twice with sterile water, adjusted to OD_600_ = 0.4 using a Tecan microplate reader (Tecan Trading AG, Switzerland). Then, 300 µL of the cell suspension was spread onto SD agar plates. After incubation for 24h at 30°C, cells were gently recovered from the agar plates with an inoculation loop and suspended in 1 mL of 0.22 µm-filtered PBS 1x in 2 mL Eppendorf tubes by pipetting up and down. The total number of cells collected from plates was evaluated by Coulter Counter (Z series, Beckman). Cells were centrifuged for 5 min at 7,000 x *g*; the supernatant was collected in a 1.5ml tube and centrifuged again at 11,000 x *g* for 5 min. Supernatants were transferred to 96-well filter plates with 0.45 µm pore size and centrifuged at 2,500 x *g* for 3 min at 4°C. The filtered supernatants were kept at −80°C for further evaluations, including the quantification of EV single particles and EV size diameter by the Flow Nanoanalyser^56^ (nanoFCM^TM^) and/or by measuring the total sterol amount by Amplex™ Red Cholesterol Assay Kit (ThermoFisher, A12216), following manufacturer’s instructions.

In the TF mutant collection^25^, each mutant is present as two or three independently obtained strains. Here, we first tested one mutant per gene. We then selected the most promising candidates producing less than 0.5-fold or more than 1.5-fold EVs compared to the WT strain. We then tested the second and third independent mutants when available in the collection. For all the mutants tested, the EV production was measured by the total amount of sterol normalized by the total number of cells collected for each strain. For the mutant strain 3.C10 (*gat1Δ*), which did not grow on SD plates, the EV production ratio was calculated based on mutant and WT growth on YPD agar plates.

### Quantification of EV particles by nanoFCM

A NanoAnalyzer (nanoFCM) instrument equipped with a 488 nm laser at 50 mW and single-photon counting avalanche photodiodes detectors (SPCM APDs) was used for the detection of the EV particles. Band-pass filters allowed for the collection of light in specific channels (488/10 nm). Light scattering and fluorescence of individual EVs were collected on single-photon counting avalanche photodiodes detectors on three channels: side scatter (SSC) - 488/10 (trigger channel). Sample fluid was focused to approximately 1.4 μm using an HPLC-grade water filter and de-gas as the sheath fluid via gravity feed. Data were generated through the Nanoanalyser Professional Suite V 2.0 software, 0.02 μm-filtered PBS was used to define the event triggering threshold. Measurements were taken over one-minute periods at a sampling pressure of approximately 1.0 kPa, modulated, and maintained by an air-based pressure module. Samples were diluted in 0.02 μm-filtered PBS as required to allow 3,000 to 13,000 counts to be recorded during this time. During data acquisition, the sample stream is completely illuminated within the central region of the focused laser beam, resulting in approximately 100% detection efficiency, which leads to accurate particle concentration measurement via single-particle enumeration. Optical alignment was tested and calibrated using fluorescent 250 nm silica nanoparticles.Further calibration measurements were taken prior to analysis using 250 nm silica nanoparticles of known concentration (for EV concentration calculation). Isolated EV samples were sized according to standard operating procedures using the proprietary 4-modal silica nanosphere cocktail generated by nanoFCM to allow for a standard curve to be generated based on the four sizes of the nanosphere populations of 68 nm, 91 nm, 113 nm, and 155 nm in diameter. Silica provides a stable and monodisperse standard with a refractive index of approximately 1.43 to 1.46, which is close to the range of refractive indices reported in the literature for EVs (*n* = 1.37 to 1.42). Using such a calibration standard enabled accurate flow cytometry size measurements, as confirmed when comparing flow cytometry with cryo-TEM results. The laser was set to 10 mW and 10% SSC decay. Data reported in the figure were handled within the nanoFCM Professional Suite v2.0 software to analyse particles between 40 nm and 155 nm.

### DNA and RNA purification and sequencing libraries preparation

*C. neoformans* KN99α, *hap2Δ,* and *gat5Δ* strains were grown in EV production condition on SD for 18h at 30°C. Cells were collected from the agar plates and suspended in 10ml 0.22µm-filtered PBS 1x. RNA extracts were prepared as previously described^57, 58^. Each condition was used to prepare biological triplicate samples. RNA-seq analysis was performed as previously described^59^. Briefly, strand-specific, paired-end cDNA libraries were prepared from 1.5 µg of total RNA by polyA selection using the TruSeq Stranded mRNA kit (Illumina) according to the manufacturer’s instructions. cDNA fragments of ∼400 bp were purified from each library and confirmed for quality by Bioanalyzer (Agilent). DNA-Seq libraries from 2.5 µg of genome DNA were prepared using the TruSeq DNA PCR-free kit (Illumina). Then, 100 bases were sequenced from both ends using an Illumina NextSeq500 instrument according to the manufacturer’s instructions (Illumina).

### Sequencing library trimming and mapping

The paired reads from the RNA-seq libraries were trimmed for low-quality reads, and Illumina TruSeq adapters were removed with Cutadapt v1.9.1 (https://doi.org/10.14806/ej.17.1.200) with the following parameters: --trim-qualities 30 –e (maximum error rate) 0.1 --times 3 --overlap 6 -- minimum-length 30. The cleaning of rRNA sequences was performed with Bowtie2 v2.3.3^60^ with default parameters; unmapped paired reads were reported using option --un-conc to identify reads that did not align with rRNA sequences. The cleaned reads from RNA-seq paired-end libraries from *C. neoformans* to the H99 reference genome (NCBI Genome Assembly GCA_000149245.3) with Tophat2 v2.0.14 ^61^ and the following parameters: minimum intron length 30; minimum intron coverage 30; minimum intron segment 30; maximum intron length 4000; maximum multihits 1; microexon search. Analysis of differential expression data was performed in DeSeq2^62^.

DNA read alignment, variant detection, and ploidy analysis were performed as previously described^63^. Illumina reads were aligned to the *C. neoformans* H99 reference genome using Minimap2 aligner v.2.9^64^ with the “-ax sr” parameter. BAM files were sorted and indexed using SAMtools^65^ version 1.9. Picard version 2.8.1 (http://broadinstitute.github.io/picard) tools were used to identify duplicate reads and assign correct read groups to BAM files. SAMtools version 1.9 and Picard version 2.8.1 were then used to filter, sort, and convert SAM files and assign read groups, and mark duplicate reads. Single-nucleotide polymorphisms (SNPs) and insertions/deletions (indels) were called using Genome Analysis Toolkit version 3.6 with ploidy=1 according to the GATK Best Practices. HaploScore parameters used to filter SNPs and indels included VariantFiltration, QD <2.0, LowQD, ReadPosRankSum<-8.0, LowRankSum, FS >60.0, HightFS, MQRankSum<-12.5, MQRankSum, MQ <40.0, LowMQ, and HaplotypeScore >13.0. To examine variations in ploidy across the genome, the sequencing depth at all positions was computed using SAMtools^65^, and then the average depth was computed for 1-kb windows across the genome. Gene ontology (GO) analyses were performed using the FungiDB database (https://fungidb.org/fungidb/app).

### Obtention of FLC-resistant *in vitro* strains and passaged derivatives

To obtain FLC-resistant *C. neoformans* strains *in vitro*, WT KN99α cells, previously grown for 24h at 30°C, were plated on drug-free YPD agar plates and incubated for 48h at 30°C. Four isolated parental colonies were used to inoculate four YPD liquid cultures grown ON at 30°C. An aliquot of the cells was stored at −80°C, and 1 ml of the culture was washed once with sterile water and adjusted to 10^5^ cells/mL. 200 µL of the cell suspension was plated on new YPD plates supplemented with FLC at 15µg/mL and incubated for five days at 30°C. Four independent FLC-resistant colonies for each initial parental culture were collected from the plates, totalizing 16 resistant isolates. Each of these 16 strains was cultured in liquid YPD drug-free medium ON before being stored at −80°C (P0 isolates). For the culture passages, susceptible parental cells and FLC-resistant derived strains were thawed on YPD agar plates (30°C) for 48h. One small loop of the cells was inoculated in 1 mL of YPD in 96 deep well plates that were incubated under rotation at 30°C. Every 48h days, 10 µL of the cell suspension was collected and inoculated in 990 µL of fresh YPD using a multichannel. The remaining cells from each passage were centrifuged at 4000 x rpm for 5 min (4°C), the supernatant was discarded, and the cell pellet was suspended in 200 µL of glycerol (40%) for keeping at −80°C. Parental strains and 96 well plates containing cells from passages 1 to 6 were stored at −80°C and further used for EV isolation and FLC-disk assays.

### Disk diffusion and spot assays

Disk diffusion assays were performed as previously described^42^, with minor adjustments. Cells were grown for 24h in YPD broth at 30°C under agitation, centrifuged, washed once in sterile PBS 1x, and diluted at 1 × 10^6^ cells/mL. 100µL of the cell suspension was spread onto YPD agar plates. A single 25 μg FLC disk (BioRad) was placed in the center of each plate. Plates were incubated at 30 °C for 48 h and 72 h and photographed individually using a PhenoBooth+ (Singer Instrument, USA) apparatus. Analysis of the disk diffusion assay was done by ImageJ or by using the diskImageR pipeline, as previously described^66^. Radius of inhibition levels, referred to as RAD throughout the manuscript, represents parameters measured at 80% drug inhibition (RAD_80_). Cells for spot assays were prepared following the same steps previously described. Cells were diluted to 1 × 10^7^ cells/mL and spotted (3 μl) in 10-fold serial dilutions onto the YPD plates containing the different drugs and compounds. The susceptibility assays were repeated a minimum of two times.

### RT-qPCR for *ERG11, AFR1-AFR3* expression, and chromosomal ploidy analysis

For the analysis of gene expression by qPCR, three independent samples were prepared for each strain (WT, *hap2Δ, gat5Δ,* and *nrg1Δ*). Cells were cultured in YPD for 18 h at 30°C, under agitation. Then, 6.4 x 10^8^ cells were transferred to 100 mL of YPD and grown for 4h, at 30°C, under agitation. After the 4 h incubation, FLC was added at a final concentration of 10 µg/mL. We skipped this step in the control. Cell cultures were incubated for 2 additional hours at 30°C under agitation. Cells were counted and washed once with sterile water, and pellets were kept at −80°C for further RNA extraction, as previously described^35^. Purified RNA was treated with DNase I recombinant, RNase-free (Roche) to eliminate residual genomic DNA. Synthesis of cDNA was performed using the Thermo Scientific Maxima First Strand cDNA Synthesis Kit for RT-qPCR, following the manufacturer’s instructions. The expression levels of *ERG11* and *AFR1-AFR3* genes were determined by qPCR using 5 µl of the appropriate dilutions of cDNA and 1 µL of each primer of interest (10 µM), listed in **Supplementary Table S4**. The reactions were performed using the SsoAdvanced Universal SYBR Green Supermix in Hard Shell qPCR plate 96-well thin wall (Bio-Rad). Amplification reactions were performed using an RT-PCR Detection System thermal cycler (Bio-Rad). The Ct values obtained in triplicates were averaged, and the number of molecules measured by standard curves and normalized to that of the housekeeping gene *ACT1* (CNAG_00483). All RT-qPCR data were from three biological replicates (three independent RNA preparations).

We used qPCR assays and primers specific to the genes CNAG_00483, CNAG_00047, and CNAG_03602 to evaluate the copy number of the chromosome 1 right harm, chromosome 1 left harm, and chromosome 2, respectively (**Supplementary Table S4**).

### Lipid analysis by thin layer chromatography

Cells collected after 24h grown on SD agar plates with or without FLC (0.6µg/mL) were suspended in a mixture of chloroform (C) and methanol (M), C/M (2:1 [vol/vol]) and incubated for 4h at room temperature. The suspension was clarified by centrifugation, and the supernatants were stored. Cell pellets were again extracted with a mixture of C and M (1:2 [vol/vol]) under the same conditions. Supernatants containing the lipid extracts were combined and dried using a rotavapor (Heidolph, Germany). The resulting cell lipid extracts were submitted to a partition system composed of C/M/0.75% KCl in water (8:4:3 [vol/vol/vol]) and vigorously mixed. The lower phase was dried under a N_2_ stream, weighed, and suspended in C to reach a concentration of 10 mg/mL of total cellular lipid extracted. To analyze cellular lipids, 50µg of total lipid was spotted into thin layer chromatography (TLC) plates (silica gel 60 F_254_; Merck, Germany). A total of 3 µL of purified Lanosterol and Ergosterol (1mg/mL) were spotted in TLC plates as running standards. The plates were developed in chambers pre-saturated for 10 min at room temperature using cyclohexane and ethyl acetate (3:2 [vol/vol] as a solvent running system. Plates were sprayed with a solution of 50 mg ferric chloride (FeCl_3_) in a mixture of 90 mL water, 5 mL acetic acid, and 5 mL sulfuric acid. After heating at 100 °C for 3–5 min, the sterol spots were identified by the appearance of a red-violet color. TLC plates were imaged, and the bands were densitometrically analysed using Image J (NIH, USA). Values were expressed as ratios between cellular lanosterol and ergosterol-specific levels.

### Statistical analysis

All statistical analyses were performed using GraphPad Prism 9 software (GraphPad Software Inc.). Data sets were tested for normal distribution using Shapiro-Wilk or Kolmogorov-Smirnov normality tests. In the cases in which the data passed the normality test, they were further analysed using the unpaired Student’s t test or ordinary one-way ANOVA. When at least one data set was nonnormally distributed, we used the nonparametric Kolmogorov-Smirnov or Kruskal-Wallis test.

### Availability and accession number

Raw data are available at Bioproject: PRJNA928602

## Acknowledgments

JR was supported by the Pasteur-Roux-Cantarini fellowship of Institut Pasteur and by FAPERJ - Fundação de Amparo à Pesquisa do Estado do Rio de Janeiro (E-26/204.456/2021). This work was supported by a CAPES COFECUB grant n°39712ZK to GJ, MLR, and LN. The work was supported by an ANR grant (Projet ResistEV AAPG2021 CE35) to GJ, IVE and AA, T.T.V.D was supported by the Pasteur - Paris University (PPU) International PhD Program. IVE is supported by a CIFAR Azrieli Global Scholar Award. We would like to thank in particular Laëtitia Viengsavanh for the help with the initial steps of TF mutant collection screening, and some members of the French Cryptococcosis Study Group: Christine Bonnal (Hôpital Bichat-Claude Bernardn Paris), Marie-Elisabeth Bougnoux (Hôpital Necker-Enfants Malades, Paris), Nathalie Bourgeois (CHU Montpellier), Sophie Cassaing (CHU Toulouse), Lilia Hasseine (CHU Nice), and Stéphane Ranque (IHU Méditerranée Infection, Marseille).

## 5) Author contributions

J.R., A.T., and F.M. performed the transcription factor mutant library screening for EV production, drug susceptibility assays, isolation of *in vitro* clinical isolates, growth passages and RT-qPCR analyses. F.M. and I.M. helped with the mutant construction. J.Y.C. and JR prepared the DNA and RNA sequencing libraries. C.M. performed the sequencing library trimming, mapping and bioinformatic analyses. P.C. and S.N. helped with the nanoflow cytometry analyses. I.V.E. advised on drug susceptibility assays and experimental design. G.P.A. performed cryo-electron microscopy analyses. L.N. and A.G.Z. performed the lipid analysis. T.T.V.D. and J.R. performed the gene complementations. A.A and M.D. selected the clinical strains to be studied. J.R. and G.J. conceived and designed the project. G.J. and M.L.R. supervised the research. J.R. and G.J. wrote the manuscript. All authors commented on the final version. All authors have read and agreed to the published version of the manuscript.

## Notes

### Competing Interest Statement

The authors have declared no competing interest.

### Summary of Updates

Minor modifications in the text Figure 12 has been deleted Figure 13 (formaly fig 14) has been improved

