## supplementary figures for "Coregulation of extracellular vesicle production and fluconazole susceptibility in *Cryptococcus neoformans*"

**Rizzo *et al*., Supplementary data**
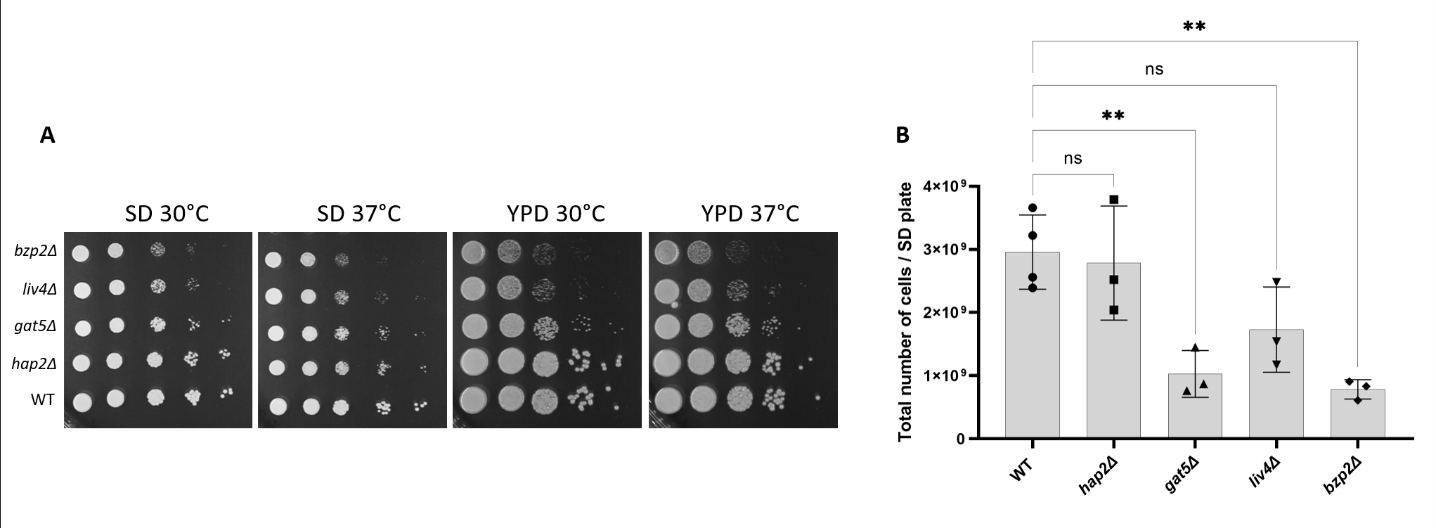

**Figure S1**: Spot assays for EV-defective mutants grown in SD and YPD agar media for 48h, at 30°C and 37°C (A). Growth analysis by counting the total number of cells grown on EV-producing conditions, SD agar plates at 30°C for 24h (B).

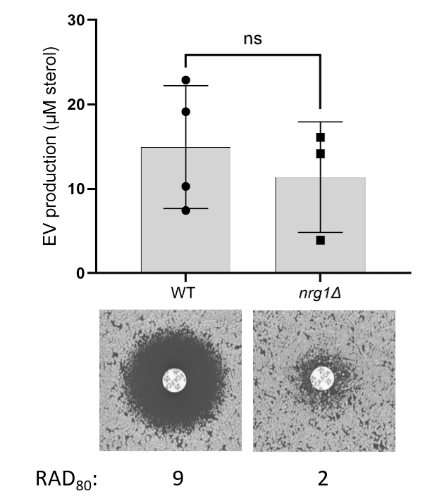

**Figure S2:** EVs production by the WT and the *nrg1Δ* mutant strains as measured by sterol concentration in the cell supernatant. Sterol concentration values are expressed per 10^9^ cells in each culture (upper panel). FLC susceptibility by disk diffusion assay (25µg/disk) and RAD analyses by diskImageR, RAD_80_ (bottom panel).

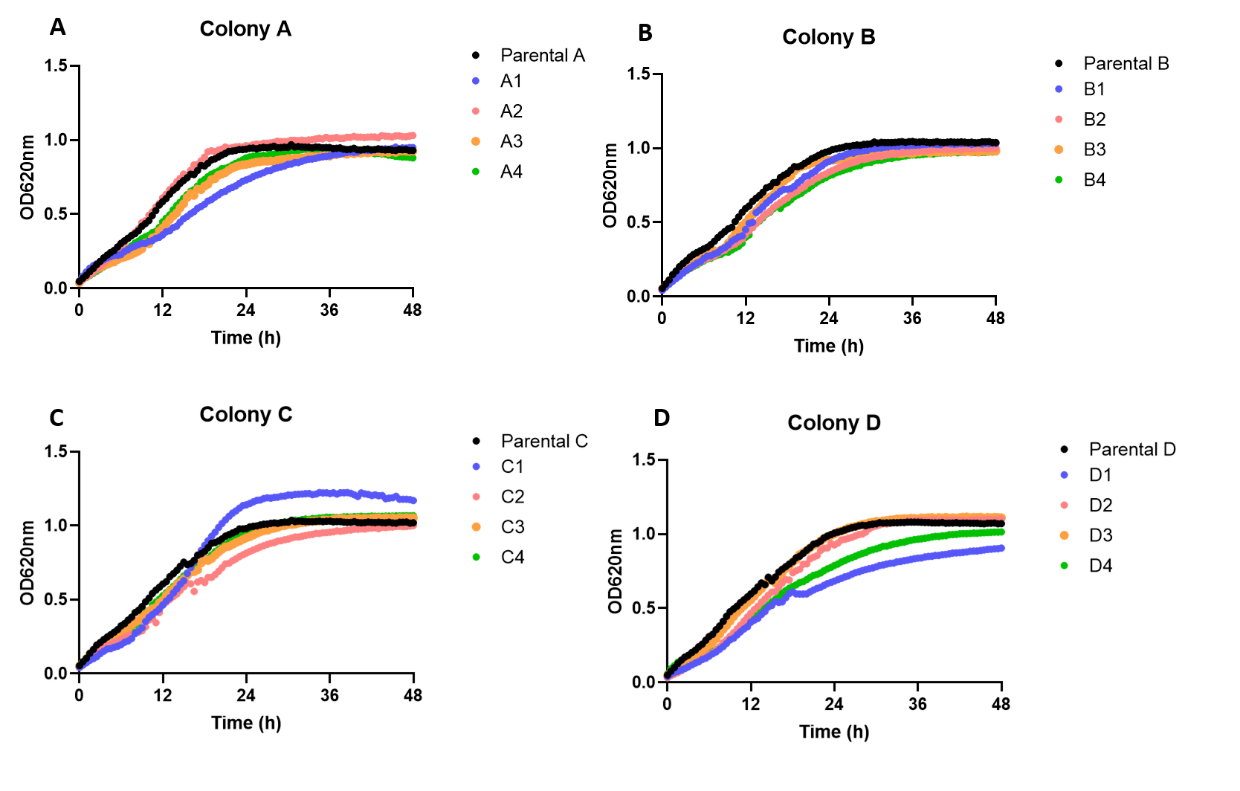

**Figure S3:** Growth analysis of parental WT strains (A, B, C, and D) and the sixteen FLC-resistant isolates derived from each parental (A1-A4, B1-B4, C1-C4, and D1-D4) incubated for 48h at 30°C under shaking on SD liquid 96-well plates. Absorbance (OD620nm) was measured every 30 minutes under a Tecan Sunrise microplate reader.

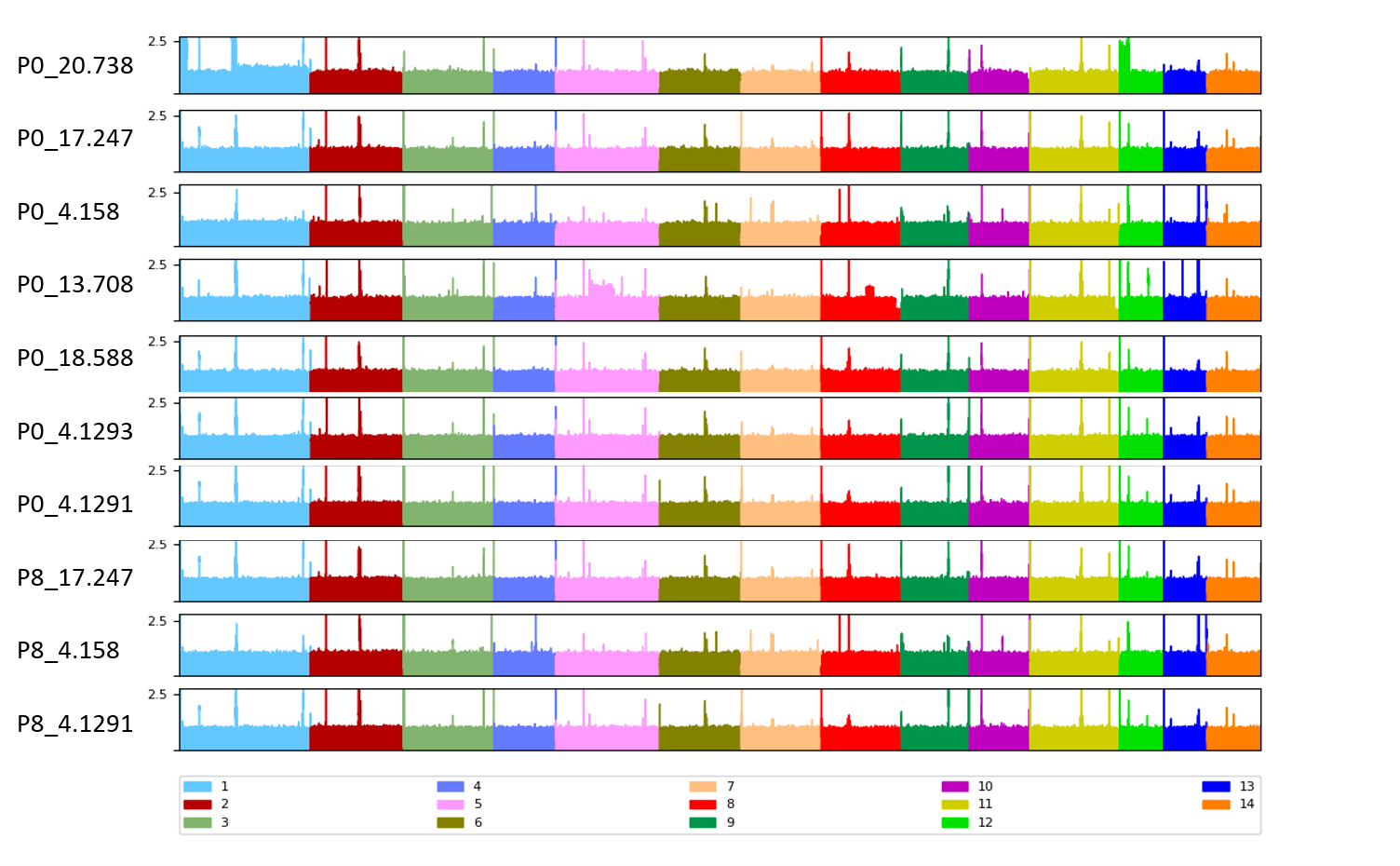

**Figure S4:** Alignment of DNA-Seq reads obtained from clinical isolates in passage 0 (P0) or passage 8 (P8).
